# Automated preparation of plasma lipids, metabolites, and proteins for LC/MS-based analysis of a high-fat diet in mice

**DOI:** 10.1101/2024.07.18.602745

**Authors:** Ngoc Vu, Tobias M Maile, Sudha Gollapudi, Aleksandr Gaun, Phillip Seitzer, Jonathon J O’Brien, Sean R Hackett, Jose Zavala-Solorio, Fiona E McAllister, Ganesh Kolumam, Rob Keyser, Bryson D Bennett

**Author notes:** To whom correspondence should be addressed Bryson D. Bennett, Director of Metabolomics Calico Life Sciences LLC 1130 Veterans Blvd, South San Francisco, CA 94080 Work. Abbreviations: Lipid classification, nomenclature and structural representation of lipids used by the LIPID MAPS; liquid-liquid extraction (LLE); cytosolic malic enzyme (ME1); Apolipoprotein C-III (ApoC-III); Apolipoprotein E (ApoE); Fibronectin (Fn1); betaine-homocysteine methyltransferase (BHMT).

## Abstract

Blood plasma is one of the most commonly analyzed and easily accessible biological samples. Here, we describe an automated liquid-liquid extraction (LLE) platform that generates accurate, precise, and reproducible samples for metabolomic, lipidomic, and proteomic analyses from a single aliquot of plasma while minimizing hands-on time and avoiding contamination from plasticware. We applied mass spectrometry to examine the metabolome, lipidome, and proteome of 90 plasma samples to determine the effects of age, time of day, and a high-fat diet in mice. From 25 μL of mouse plasma, we identified 907 lipid species from 16 different lipid classes and subclasses, 233 polar metabolites, and 344 proteins. We found that the high-fat diet induced only mild changes in the polar metabolome, upregulated Apolipoproteins, and induced substantial shifts in the lipidome, including a significant increase in arachidonic acid (AA) and a decrease in eicosapentaenoic acid (EPA) content across all lipid classes.

## 1. Introduction

Mass spectrometry-based metabolomics, lipidomics, and proteomics have become essential parts of systems biology research, offering insights into biological conditions across many *in vitro* and *in vivo* systems (1–7). Due to differences in hydrophobicity and molecular weight, metabolomics, lipidomics, and proteomics generally require separate methods for extraction and enrichment of the targeted analyte class prior to analysis (8–10). Blood sample volume can often be limited in work with small organisms like mice, restricting possible analyses. Recently, the interest in preparing samples for multiple analyses has grown due to the desire to more fully characterize and generate a better understanding of complex biological systems (11–13). The ability to prepare a single sample for multiple data analysis modalities opens the possibility of multiomic analysis on small sample volumes.

The methyl tert-butyl ether (MTBE)-methanol-water liquid-liquid extraction (LLE) method for lipid extraction developed by Matyash et al. (14–16) has been demonstrated to have similar performance to the chloroform-based method developed by Folch (17), but with the advantages of avoiding chlorinated solvents and locating the protein pellet at the bottom of the vessel, making it a popular alternative to chloroform-based extractions for lipidomics. There are several reports of metabolomics analysis on the aqueous phase of this preparation in conjunction with lipidomics analysis of the organic phase (18–23).

An automated MTBE-LLE method for the preparation of lipids prior to mass spectrometry analysis was previously reported (24, 25), as was one for the Butanol:Methanol method (BUME) (15, 16). However, to date, no reports have been made of an automated method to examine both metabolomics and lipidomics separately from the same sample. Previous reports describing a single sample preparation protocol for proteomics, lipidomics, and metabolomics analysis were not automated, limiting throughput (26–28).

To enable multiomic analysis of large sample sets, we optimized a solvent system and consumables for an automated liquid-liquid extraction system that enables lipidomic, metabolomic, and proteomic analyses from a single aliquot of blood plasma. This method provides separate fractions for lipidomics and metabolomics, requiring only offline sample evaporation and resuspension. The protein pellet remains in the LLE vessel and is compatible with many downstream proteomics preparation workflows. In this work, we used the recently published, high-throughput AutoMP3 method (29).

To demonstrate the utility of our workflow and its ability to generate high-quality data from a large number of samples, we prepared and analyzed 90 mouse plasma samples, investigating the effect of age and diet on the metabolomics, lipidomic, and proteomic profiles of mouse plasma. Our study revealed differences in metabolites, lipids, and proteins between adult and adolescent mice; and profound changes induced by a high-fat diet, especially in the lipid acyl-group composition and abundance of lipoproteins.

## 2. Materials and methods

### Chemicals, Reagents and Consumables

LC-MS-grade isopropanol, methanol, acetonitrile, and MTBE were purchased from Honeywell (Charlotte, NC, USA). Ammonium formate, ammonium bicarbonate, and formic acid were purchased from Sigma Aldrich (St. Louis, MO, USA). Lipid standards, including Splash Lipidomix (P/N 330707), EquiSplash (P/N 330731), LighSplash (P/N 330732), d31-LPC 16:0 (P/N 860397), d70-PE(18:0/18:0) (P/N 860373), d54-PA(14:0/14:0) (P/N 860450) were purchased from Avanti Polar Lipids (Alabaster, AL, USA). d35-FA 18:0 (P/N 9003318) was purchased from Cayman Chemical (Ann Arbor, MI, USA), and d4-lysine (P/N 616192), d5-phenylalanine (P/N 615870), d4-succinate (P/N 293075), L-tyrosine-(phenyl-d4) (P/N 489808), L-arginine-15N4 hydrochloride (P/N 600113), d5-benzoate (P/N 586331) were purchased from Sigma Aldrich (St. Louis, MO, USA). Metabolomics QC kit and associated light standards were purchased from Cambridge Isotope Laboratories (Tewksbury, MA, USA).

Tandem mass tag (TMTpro) 16-plex isobaric reagents were obtained from Thermo Fisher Scientific (Rockford, IL, USA). Modified Trypsin and LysC mix were obtained from Promega Corporation (Madison, WI, USA). Complete protease inhibitor tablets were obtained from Roche. Thermo Fisher Scientific Pierce Bicinchoninic Acid (BCA) Assay (P/N 23228), GE Healthcare SpeedBead Magnetic Carboxylate Beads Hydrophilic and Hydrophobic forms (P/N 65152105050250, 45152105050250).

PCR Strip Caps (P/N, 321-11-071) 0.2 mL Clear, For Real Time PCR, flat top; Platemax UltraClear Sealing Film (P/N UC-500); Axygen® AxyMats™ 96 Round Well Compression Mat for PCR Microplates, Non-sterile (P/N CM-96-RD); PCR Tubes 0.2 mL Maxymum Recovery, Thin Wall, Clear (P/N 321-02-501); 2 mL 96-well deep well plate (P/N P-2ML-SQ-C) were obtained from Axygen (Union City, CA, USA). Eppendorf 2.0 mL Protein LoBind tubes (Catalog#022431102), Eppendorf 1.5 mL Protein LoBind tubes (Catalog#02243108), and Alpaqua 96-well neodymium magnet (Cat#A000400), Eppendorf 150 µL DNA LoBind 96-well PCR plates (P/N 0030129512), Eppendorf 1 mL Protein LoBind 96-well plates (P/N 951033308) were purchased from Eppendorf (Enfield, CT, USA). Corning plate lids (P/N CLS3935-50EA) from Corning (Glendale, AZ, USA). Hamilton 60 mL reagent reservoir (P/N 56694-01); Filtered, Conductive Hamilton VantageTM Tips – 1000 µL (P/N 235940), 300 µL (P/N 235938), 50 µL (P/N 235979) were from Hamilton (Cary, NC, USA). Unless otherwise stated, all other chemicals were purchased from Sigma-Aldrich (St. Louis, MO, USA).

### Blood Plasma Samples

All animal protocols in the USA were approved by the Mayo Clinic IACUC and the Calico Life Sciences IACUC. The studies were conducted under the NIH Guide for the Care and Use of Laboratory Animals (protocol # A00003888). Ten male C57BL/6 mice (Jackson Laboratory, USA) were selected for each of three groups, to minimize variance in body weight, and eliminate animals that had fighting wounds. Groups of 8 to 12-week-old (young) and 24 to 28-week-old (adult) mice were fed a standard lab chow diet (Chow diet 5LOD, LabDiet), and a second group of 24 to 28-week-old mice fed 10 weeks of a high-fat diet (HFD) containing 60% fat (D12492, Research Diets Inc.). They were housed in standard cages at room temperature and constant humidity, with a 12-hour light-dark cycle. Three blood plasma samples were obtained from each mouse via retro-orbital bleeding. Each sample was obtained on a different day and at a different time of day: 10 AM (time point 1-Z10), 10 PM (time point 2-Z22), and 3 PM (time point 3-Z15). Blood was collected into BD microtainer tubes (REF 365974) with K2E (K2EDTA), mixed briefly, and stored on ice. All samples were centrifuged at 7000 RCF for 4 mins at 4°C to separate blood plasma. Plasma was aliquoted in 50 µL increments, immediately frozen on dry ice, and stored at −80C for later analysis.

### Automation/Robotics and Consumables

Automated LLE extraction was performed on a PAL DHR Dual Head system from Trajan Scientific (Morrisville, NC, USA) equipped with modules as described in Figure 1. Customized parts for the robot included: reagent chiller assembly, buffer holder with chiller, customed aluminum vial racks, nitrogen gas line, and custom enclosure.

**Figure 1:**
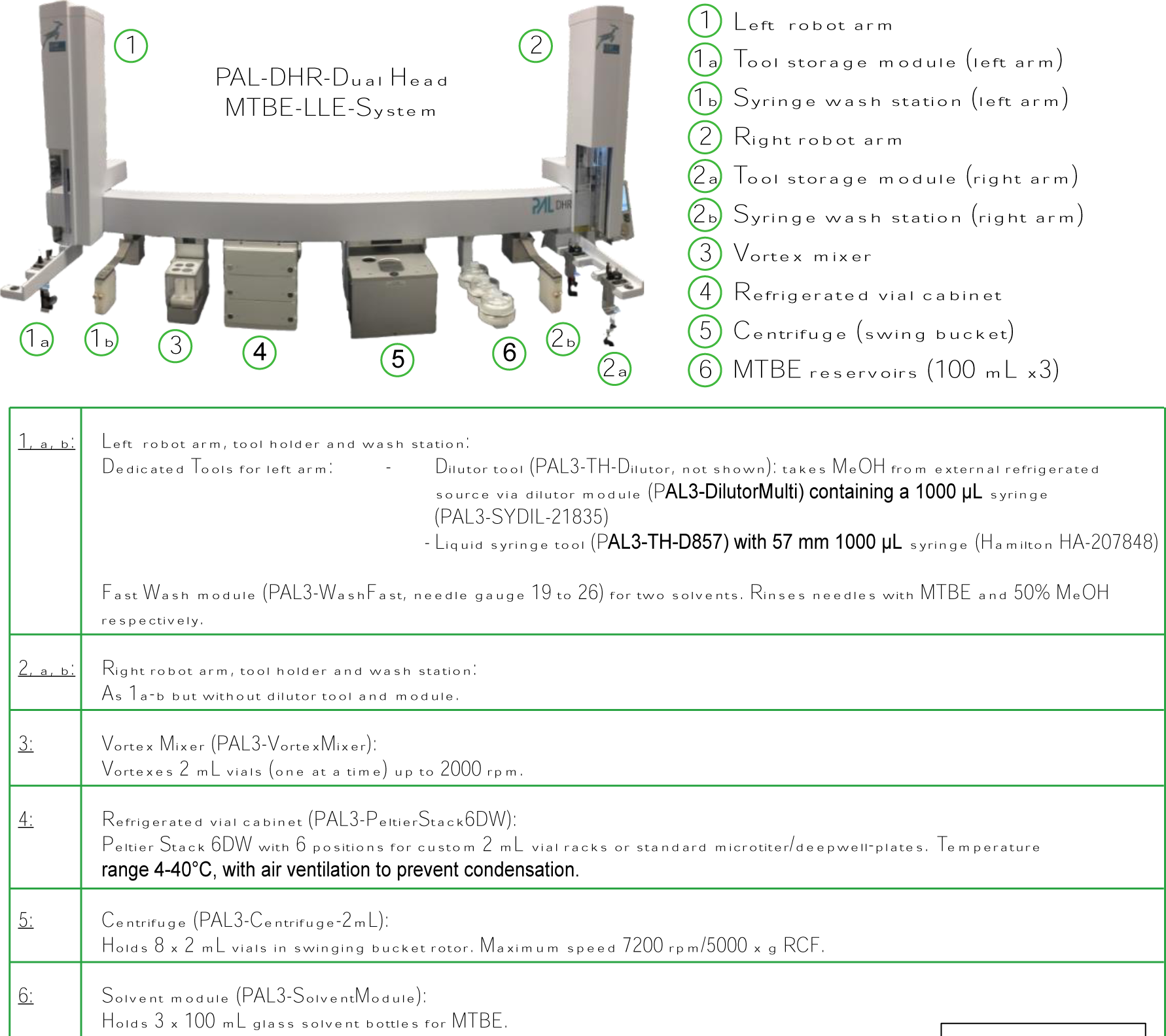
PAL-DHR diagram and description of customized parts. Plasma aliquots were stored in the 4℃ cabinet (#4). PAL-DHR used its two arms (#1, #2) for liquid transferring and relocating glass vials with magnet caps. Briefly, PAL added the extraction buffers to samples, then vortexed (#3) and centrifuged (#5) to collect lipid and metabolite fractions, which were subsequently stored in the second and third level of the refrigerated vial cabinet. Details of the automated protocol can be found in the Method section.

### Manual Liquid-Liquid Extraction

Manual LLEs were performed using a modified version of Matyash procedure (14). Briefly, a mixture of lipid standards (volumes as described in results) was extracted by the addition of 50% methanol (pre-cooled to −20°C), followed by vortexing for 30 sec and incubated on ice for 10 min. The same volume of room temperature MTBE was then added, and the samples were vortexed 30 sec before centrifugation at 3000 x g for 5 min at 4°C for complete phase-separation. The upper organic phase was aspirated with a Hamilton glass syringe until the meniscus of the two phases was reached and transferred to a new ice-cold glass sample vial. The remaining sample volume was subjected to a second extraction using MTBE or other solvent mixtures, as described in the results section. The second organic phase was removed and combined with the first organic phase extraction. All sample transfers involving MTBE-containing solutions were performed with glass syringes rinsed twice with 100% MTBE then MeOH, between preparation steps. Following both MTBE extractions, the lower aqueous phase extract was pipetted off with the necessary care not to pick up the remaining upper phase or disturb the protein pellet. The fraction was subsequently transferred into a separate ice-cold sample vial. Samples were evaporated to dryness under nitrogen at 4°C and then stored at −80°C.

### Automated Liquid-Liquid Extraction

EDTA-plasma aliquots of 25 μL were resuspended in glass vials (SureStop 2 mL vials #C5000-1W with National Scientific custom magnetic caps #CPSH0003 from Thermo Scientific) to a final concentration of 50% methanol, containing 10 μL of a mixture 1 µg/mL of deuterated tyrosine and deuterated alanine, 250 ng/mL of deuterated benzoate (Sigma Aldrich) in water, as well as 10 µL Splash Lipidomix mass spec standard (Avanti Polar Lipids) as internal standards in methanol, generating sample extraction volumes of 70µL in the source sample vials. The robot performed LLE as per the script provided in SI document.

Briefly, 50% methanol (pre-cooled to −20°C) was added via the PAL-DHR system from an external reservoir to a final volume of 800 µL, followed by vortexing for 70 sec at 2000 rpm. Samples were then automatically extracted in sets of 8 (or an even number of fewer samples for the last batch of the sample set depending on the total sample number), with all subsequent liquid movements performed using 1 mL Hamilton Syringes (Hamilton HA-207848). Lipid extraction was performed via the addition of 750 µL of MTBE to each sample vial followed by vortex mixing for 30 sec. The sample set was then centrifuged for one min at 1400g for complete phase separation, and 520 µL of the upper (organic) phase was transferred to a lipidomics collection vial (National Scientific #CPSH0008, Thermo Scientific). The second extraction was performed via addition of 600 µL of MTBE to the source vial. After repeating the mixing and centrifugation steps as described above, 860 µL of the upper (organic) phase was transferred to the lipidomics collection vial. Finally, 425 µL of the lower aqueous phase was transferred to the metabolomics collection sample vials. Throughout the procedure, sample vials, and destination vials for both the organic and aqueous fractions were stored in cold blocks chilled to 4°C in the vial cabinet (#5 in Fig.1). The vial cabinet was constantly purged with nitrogen gas throughout the automated LLE procedure. Syringes were washed between each sample movement using MTBE for organic phase transfers, and 50/50 v/v/ methanol/water for aqueous phase transfers.

### Automated Proteomics Sample Preparation

Following the LLE procedure above, the sample vial contained approximately 100 µL of the upper organic phase and 150 µL of lower aqueous phase and the protein pellet. Samples were stored at −80°C until the day of preparation at which time they were evaporated to dryness under nitrogen, then diluted 10x in protein lysis buffer (3% SDS, 50 mM HEPES pH 8.5, 75 mM NaCl + Roche complete cocktail of protease inhibitors). Samples were processed using the AutoMP3 workflow, as previously described (31) Briefly, proteins were reduced with DTT, alkylated with IAA, and cleaned up with SeraMag beads prior to protein digestion. Digestion was performed using LysC & Trypsin (Promega, 1:20 protease to protein ratio) for 1 hour (37°C, 1000 rpm). Peptides were labeled with TMTpro as 16-plexes with one bridge channel per plex. The bridge sample was created by combining a portion of all samples to make a pooled control, and labeled using the 134N TMT tag. Samples were quenched and desalted prior to LCMS analysis through the combine-mix-split strategy and peptide-level SeraMag bead cleanup steps. Proteins and peptides were quantified using a BCA assay (Pierce).

### LC-MS Analysis of Polar Metabolites

Analysis of polar metabolites was performed on a Vanquish HPLC coupled to a Q Exactive Plus mass spectrometer (Thermo Scientific). Dried aqueous phase extracts were resuspended in 100 µL of water containing 1 µg/mL of deuterated lysine and deuterated phenylalanine and 250 ng/mL of deuterated succinate (Sigma Aldrich) as internal standards. Samples were centrifuged at 18000 x g for 5 minutes, and the supernatant was moved to HPLC vials.

Metabolites were analyzed in negative ionization mode using a reverse phase ion-pairing chromatographic method with an Agilent Extend C18 RRHD column, 1.8 μm particle size, 80 Å, 2.1 × 150 mm. Mobile phase A was 10 mM tributylamine, 15 mM acetic acid in 97:3 water:methanol pH 4.95; mobile phase B was methanol. The flow rate was 200 μL/min and the gradient was t = −4, 0% B; t = 0, 0% B; t = 5, 20% B; t = 7.5, 20% B; t = 13, 55% B; t = 15, 95% B; t = 18.5, 95% B; t = 19, 0% B; t = 22, 0% B. The mass spectrometer was operated in negative ion mode using data-dependent acquisition (DDA) mode with the following parameters: resolution = 70,000, AGC target = 1.00E + 06, maximum IT (ms) = 100, scan range = 70–1050. The MS2 parameters were as follows: resolution = 17,500, AGC target = 1.00E + 05, maximum IT (ms) = 50, loop count = 6, isolation window (m/z) = 1, (N)CE = 20, 50, 100; underfill ratio = 1.0 0%, Apex trigger(s) = 3–12, dynamic exclusion(s) = 20. The injection volume was 5 µL.

For positive ion mode, the sample resuspension described above was diluted 1:4, v/v with MeCN and centrifuged at 18000 x g for 5 min, and 30 µL of the supernatant was moved to HPLC vials without disturbing the precipitate. Metabolites were analyzed in positive ionization mode via HILIC chromatography using a SeQuant® ZIC®-pHILIC column, 5 μm particle size, 200 Å, 150 × 2.1 mm.

Mobile phase A was 20 mM ammonium carbonate in water (pH 9.2); mobile phase B was acetonitrile. The flow rate was 150 μL/min and the gradient was t = −6, 80% B; t = 0, 80% B; t = 2.5, 73% B; t = 5, 65% B; t = 7.5, 57% B; t = 10, 50% B; t = 15, 35% B; t = 20; 20% B; t = 22, 15% B; t = 22.5, 80% B; t = 24; 80% B. The mass spectrometer was operated in positive ion mode using data-dependent acquisition (DDA) mode with the following parameters: resolution = 70,000, AGC target = 3.00E + 06, maximum IT (ms) = 100, scan range = 70–1050. The MS^2^ parameters were as follows: resolution = 17,500, AGC target = 1.00E + 05, maximum IT (ms) = 50, loop count = 6, isolation window (m/z) = 1, (N)CE = 20, 40, 80; underfill ratio = 1.00%, Apex trigger(s) = 3–10, dynamic exclusion(s) = 25. The injection volume was 5 µL.

### LC-MS Analysis of Lipids

Detection of lipids was performed on a Vanquish HPLC coupled to a Q Exactive Plus mass spectrometer (Thermo Scientific). Dried organic phase extracts were resuspended in 100 µL of (2:1:1,v/v/v) ButOH:MeOH:H_2_O including 500 ng/mL d70-PE(18:0,18:0), 2.5 µg/mL d31-LPC(16:0), 700 ng/mL d54-PA(14:0,14:0) and 500 ng/mL d35-FA(18:0) as deuterated LCMS lipid standards to monitor instrument performance. Separation of lipid compounds in both positive and negative ion mode was achieved by reverse-phase liquid chromatography on an Accucore C30 column (250 x 2.1 mm, 2.6 µm particle size, Thermo Scientific). Mobile phase A was 20 mM ammonium formate in 60:40 meCN:H_2_O, with 0.25 µM medronic acid and mobile phase B was 20 mM ammonium formate in 90:10 IPA:meCN, with 0.25 µM medronic acid. The gradient was: t= −7, 30% B; t=7, 43% B; t=12, 65% B, t = 30, 70% B; t=31, 88% B; t=51, 95% B, t=53, 100% B t=55, 100% B; t=55.1, 30% B, t=60, 30% B for a total run time of 67 min per injection. The flow rate was 200 µL/min, the injection volume was 5 μl, the column temperature was 35°C. Parameters for the full scan (MS^1^) were: 140,000 resolution, AGC target of 3e6, IT of 100 ms and a scan range of 200 to 2000 m/z. The MS^2^ parameters were: 17,500 resolution, a loop count of 8, and an AGC target of 3e6, IT of 150 ms, isolation window of 1 m/z and underfill ratio of 1%. Dynamic exclusion was set at 15 sec with an apex trigger from 5 to 30 sec. Stepped collision energies for fragmentation were set to 20, 30 and 40% NCE.

### LCMS Analysis of Peptides

Peptides from each combined 16-plex were resuspended in 95/5 v/v water/acetonitrile with 5% formic acid at a concentration of approximately 0.33 µg/µL and 3 µL was injected (1 µg) for analysis. Peptides were separated by reverse phase on a C18 Aurora microcapillary column (75 μm x 25 cm, C18 resin, 1.6 μm, 120 Å, #AUR2-25075C18A) (IonOpticks) on a Ultimate3000 LC (Thermo Scientifics) coupled to an Orbitrap Eclipse mass spectrometer. The total LC-MS run length for each sample was 185 min including a 165 min gradient from 8 to 30% ACN in 0.125% formic acid. The flow rate was 300 nL/min, and the column was heated at 60° C. Data was collected utilizing DDA with RTS and FAIMS Pro. We used four different FAIMS Pro compensation voltages: −40, −50, −60, and −70 Volts. Each of the four experiments had a 1.25 seconds cycle time. A high-resolution MS1 scan in the Orbitrap (m/z range 400-1,600, 120k resolution, standard AGC Target, “Auto” max injection time, ion funnel RF of 30%) was collected, from which the top 10 precursors were selected for MS2, followed by SPS MS3 analysis. For MS2 spectra, (30) ions were isolated with the quadrupole mass filter using a 0.7 m/z isolation window.

The MS2 product ion population was analyzed in the ion trap (CID with normalized collision energy 35%, AGC 1 x 10^4^, custom max injection time of 35 ms). The minimum Xcorr needed to pass the RTS search was set to 2, the minimum dCn was set to 0.1, and a maximum of two missed cleavages was allowed. The MS3 scan was analyzed in the Orbitrap (50k resolution and with a scan range of 100-500 m/z, HCD with fixed normalized collision energy of 45%, AGC 1 x 10^5^, max injection time 200 ms). Up to ten fragment ions from each MS2 spectrum were selected for MS3 analysis using SPS.

### Data Processing

For metabolomics and lipidomics data, raw files were converted to mzML format using Proteowizard Ver 3 (30). Compound identifications and peak grouping were performed using the <u>OpenCLaM</u> R package (https://github.com/calico/open_clam). Peaks were matched within 10 ppm for the precursor mass and 20 ppm for fragment masses. Fragmentation and retention times were compared to an in-house library generated from authentic standards for metabolomics, and fragmentation was compared to in-house generated in-silico library for lipidomics (31). Following automated annotation, the data set was manually inspected, validated, and some identifications were reassigned using MAVEN2 (31). Each metabolite feature was log_2_ transformed and normalized to the mean of the young-chow mice using <u>claman</u> R package (https://github.com/calico/claman). Lipidomics data were normalized to the isotopically labeled internal standard of the same class from Splash Lipidomix (Avanti Polar Lipids). Absolute quantification of each lipid species was calculated based on the ratio of the peak area top between feature and internal standard. Annotation was according to the LIPID MAPS classification, nomenclature, and shorthand notation guidelines (32).

Proteomics mass spectrometry data were processed using Masspike software (GFY Core) Version 3.8 (33). Raw files were converted to mzXML files using ReAdW and searched against either a mouse Uniprot database (downloaded on January 8th, 2020) in forward and reverse orientations using the Sequest algorithm (34). Database searching matched MS/MS spectra with fully tryptic peptides from this dataset with a precursor ion tolerance of 20 ppm and a product ion tolerance of 0.6 Da. Carbamidomethylation of cysteine residues (+57.02 Da) was set as static modification, and TMTpro tags of peptide N-termini and lysines (+304.20 Da) were set as static modifications as well. Oxidation of methionine (+15.99 Da) was set as a variable modification. Linear discriminant analysis was used to filter peptide spectral matches to a 1 % FDR (false discovery rate) as described previously (33). Non-unique peptides that matched to multiple proteins were assigned to proteins that contained the largest number of matched redundant peptide sequences using the principle of Occam’s razor (33). Quantification of TMT reporter ion intensities was performed by extracting the most intense ion within a 0.003 m/z window at the predicted *m/z* value for each reporter ion. Peptide intensities and signal-to-noise ratios were exported from in-house software and analyzed using msTrawler software package (35).

### Statistics

For the comparison on different ages and diets, statistical analysis was performed using the claman R package for metabolomics and lipidomics, and the msTrawler R package for proteomics (35). Each metabolite/lipid/protein were modeled separately with a mixed-effect model that included fixed effects for age, diet, and blood draw batch, and a random effect for each individual mouse using the formula: 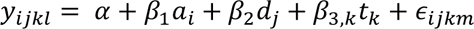, where *y*_*ijkl*_ are the log_2_ reporter ion intensities for metabolites, and log_2_ concentration for lipids, observed from the feature being modeled. Observations indexed by i = 1 are from young mice, and i = 2 from the old mice (α_1_= 0 and α_2_= 1), so that β_1_ is the expected change in log_2_ abundance between young and old mice. j =1 or 2 indexes the diet category so that *d*_1_= 0 for mice on the chow diet and *d*_2_= 1 for mice on the high-fat diet. Accordingly, β_2_ represents the expected change in log_2_ abundance when moving from a chow to a high-fat diet. k = 1, 2 or 3 indexes the three sampling times, Z10, Z22 or Z15. The first time is the reference *t*_1_ = β_3,1_ = 0 and *t*_2_ and *t*_3_ equal 1 when the sampling time were at time Z22 or Z15 respectively and take values of zero otherwise so that β_3,2_ and β_3,3_ represent expected changes in relative abundance due to blood draw batch effects. *b*_*m*_ ∼ *N*(0, τ^2^) is a random intercept for each mouse that creates a correlation structure in the model between all repeat measurements taken on the same mouse. τ^2^ is the variance of relative abundance between animals.

For proteomics, we included a term to account for the measurement error from sample processing. Each *b*_*m*_ is mutually independent of one another and of the residual variance, ε_*ijkm*_ ∼ *N*(0, *w*_*ijkm*_σ^2^). *w*_*ijkm*_ is the inverse of the ion count for each scan, and bridge channels were incorporated into the model as previously described (35). False Discovery Rate was controlled on a term-by-term basis using the qvalue R package (36). Significant changes were reported for q values < 0.01.

To assess the overall impact of age, diet, and blood draw time point in shaping variability in the metabolome and lipidome, we assessed how much of the total variability in the dataset is explained by each factor. To do this we fit a three-way ANOVA using the formula 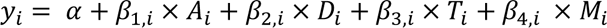 to each feature using the R aov() function. This decomposed the variance of each feature (described by the total sum of square (TSS); i.e., Var(y)*N) into variance explained by each experimental factor (explained sum of squares (ESS)) and unexplained variance (residual sum of squares (RSS)). The ratio of ESS to TSS provides a measure of the relative effect size of each variability. To provide an estimate of this ratio at a dataset-level, we summed the ESS of each variable and the TSS across all features and then took the ratio of dataset-level ESS to TSS as a summary of total variance explained by each variable.

For lipid enrichment analysis, we ranked all identified species using the estimated values calculated from the regression model. Identified lipid species were separately classified into lipid classes, and acyl chain composition, which were used as the groups of which to determine enrichment. Then, lipid class and acyl enrichment were performed on the ranked species using R package FGSEA in both directions (37–41). We report the -log_10_ Benjamini-Hochberg corrected p-value and use the normalized enrichment score (NES) to assign the direction of the change.

## 3. Results

### 3.1 Automation of MTBE-LLE extraction

Extractables and leachables from plastics and adsorption of hydrophobic materials onto plasticware are significant concerns in the fields of lipidomics and lipid biochemistry as they can mask differential abundance of lipids between samples (42–44). To determine the effects of plasticware on the results obtained during MTBE-based extraction, we performed an untargeted analysis of features from LC-MS lipidomics analysis of 10 µL of aliquots mouse plasma extracted in either 1.5 mL glass vials or low-retention polypropylene microcentrifuge tubes. We found many features with higher abundance in the samples extracted in polypropylene, most of which did not match known lipids. We putatively identified some of these compounds as chemicals used in polymer production, including triton and polybutylenes (Figure S1). We identified monoglycerides and fatty acids which were extracted from the plastic, including palmitic acid, stearic acid, palmitoylmonoglyceride, and stearoylmonoglyceride.

To avoid the contaminants observed with polypropylene vessels in the automated system, we implemented an LLE method on a PAL-DUAL system (Fig.1), a dual-head robotic platform capable of automatic tool changes that use glass syringes for liquid transfers. We configured the system with cooled blocks for the 2 mL glass vials (4℃), chilled solvents (4℃), and constantly purged the vial cabinet with nitrogen to minimize oxidation. The system consisted of a vial cabinet, vortex mixer, centrifuge, and syringe wash stations, allowing for the complete automation of the entire LLE as shown in Fig.1. We enclosed the instrument to allow any solvent vapor to be ventilated directly to the building exhaust system.

Initially, we attempted to implement the LLE using the solvent ratios described in Matyash et al: a first extraction with a solvent composition of MTBE/MeOH/H_2_O (10/3/2.5, v/v/v) and a second extraction composed of MTBE/MeOH/H_2_O (2/1/1, v/v/v), an approximation of the upper phase composition of the first extract. However, we found that this 2/1/1 mixture separated into two phases over time, likely due to small amounts of evaporation changing the solvent composition. To enable automation, a reservoir with a single phase was preferable, so we examined the impact of either eliminating the second extraction step or using 100% MTBE, while leaving the first extraction unchanged. Without the second extraction step, we recovered significantly less phospholipid and lysophospholipids, but we found similar recoveries using MTBE/MeOH/H2O (2:1:1, v/v/v) or 100% MTBE (Fig. S2A and Fig. S2B). We settled on a final protocol consisting of a first extraction of MTBE/MeOH/H2O (2/1/1, v/v/v) solvent ratio and a second extraction of 100% MTBE addition to the remaining aqueous phase.

In tests of the automated LLE, we found a series of high-intensity contaminants during LC-MS analysis despite the careful solvent handling to avoid exposure to plastics. We identified the contaminants as polysiloxanes (Fig S3A), which organic solvents may extract from silicone. We found that the repeated penetration of the septa during preparation fractured the PTFE layer of the cap, exposing the silicone to MTBE during vortex mixing. We compared caps made with PTFE-silicone, PTFE-red rubber, or a plain PTFE disc and found a 10-fold reduction in peak size with the PTFE-red rubber septum and more than a 100-fold reduction in peak intensity with the PTFE disc. Hence, we moved forward with the PTFE disc for this method (Figure S3B).

However, when performing automated preparations with PTFE-disc closures, we found that the standard 22 gauge, type 3 needle (a blunt needle with the port at the tip) would occasionally core the PTFE disc, causing either clogging of the syringe or debris in the sample. To address this problem, we tested two other needle types: the narrower, conical-tip AS needle with a bottom port and the conical-tip type 5 needle with a side port. We found that the AS needle clogged less, but did still occasionally clog, hence we chose the type 5 needle, which is more commonly used for gas chromatography injections, as it showed consistent performance.

In order to allow for slight differences in samples and ensure that the incorrect phase or protein precipitate would not contaminate the destination vials, we chose conservative volumes to be transferred out of the sample vial, leaving approximately 100 uL of the organic phase and 150 uL for the aqueous phase behind with the protein pellet, giving us 74% of the aqueous phase volume (425 of 575 uL) and 93% of the total organic phases (1380 of 1480 μL).

### 3.2 Validation of the automated MTBE-LLE method

We examined the fractional recovery of 10 polar metabolites, and 13 common lipid classes using pure chemical standards. We recovered 50% to 86% of lipids, with the lowest recovery observed for the most hydrophobic lipid classes, triglycerides (TG) and cholesterol esters (CE), consistent with previous reports on the extraction efficiencies of methyl tert-butyl ether (MTBE)-based methods (45–47). For polar metabolites, the recovery ranged from 30% to 78%, exhibiting a strong correlation with the hydrophobicity of the metabolite (Fig. 2A). The automated method yielded results similar to those obtained through manual extraction for metabolites and slightly lower yields for lipids (Figure S4A and S4B).

**Figure 2:**
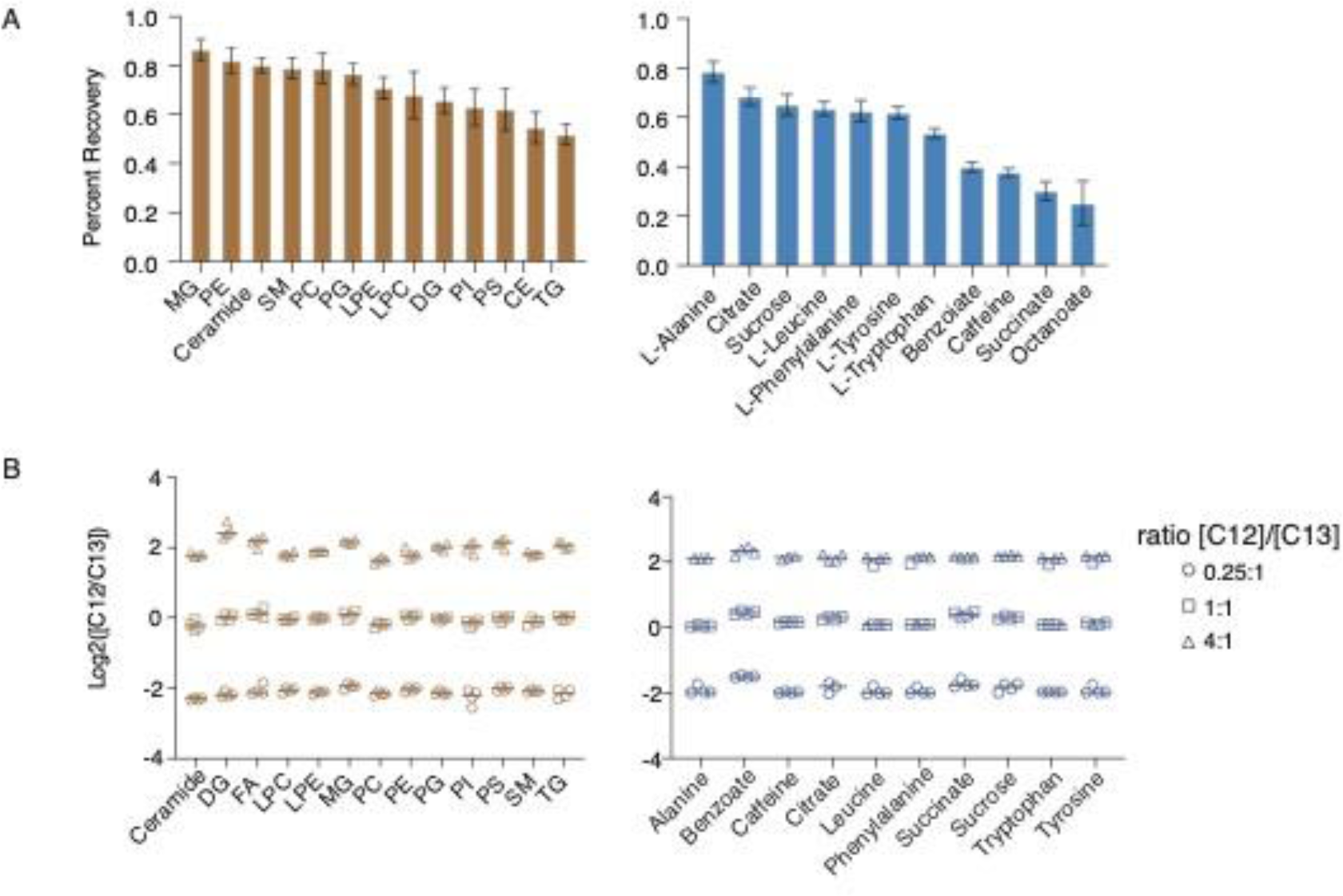
Accuracy and Precision of extraction recovery from the automated MTBE-LLE-PAL system. A) Extraction efficiency reported by MTBE-LLE performed by PAL system. Log2 ratios reflect PAL-LLE prepared isotopically labeled internal standards relative to equimolar unlabeled standards added post-LLE (PAL, n=4). Lipid recovery in the organic layer (brown); Metabolite recovery in the aqueous layer (blue). B) Accuracy and precision of the automated MTBE-PAL. Standard mixtures containing isotopically labeled standards (heavy) and their associated C12 standards (light) were prepared at the ratio (1:0.25, 1:1, 1:4, v/v, (n=3), followed by LLE separation. Ratios were normalized to controls that did not undergo the LLE.

Next, we evaluated the accuracy and precision of the automated sample preparation. We prepared three concentrations of unlabeled standards, spanning a 16-fold concentration change, with unvarying isotopically labeled internal standards (Fig. 2B). We found high linearity with R^2^ greater than 0.96 for all compounds, and a median CV of 11%, a minimum (minimum 6%, maximum 23%) for repeated measurement of the labeled isotopic standards at the same concentration.

To assess the batch-to-batch reproducibility of the system, we extracted varying volumes of aliquots from the same plasma in over two separate distinct weeks, and then acquired LC-MS data in a single batch. We performed a regression on the mean peak area (n=3) of lipids and metabolites identified from each batch. High correlations were determined in both fractions, with R^2^ = 0.998 for metabolites and R^2^ = 0.996 for lipids (Fig. S4C, Fig. S4D).

Finally, we determined the carry-over of the automated method by sequentially preparing plasma standards and water blanks. We found minimal carry-over, with less than 2% of the signal of the ten most abundant plasma metabolites and the ten most abundant lipids per lipid class found in the blank (Fig. S4E).

### 3.3 Determining the effects of age and a high-fat diet on mouse plasma composition

To demonstrate the utility of the method described above in analyzing biological samples, we prepared 90 mouse plasma aliquots of 25 μL each. The samples comprised three groups of n=10 mice: 8-week-old mice (young-chow) and 24-week-old mice (adult-chow) fed a standard chow diet, and 24-week-old mice (adult-HFD) fed a high-fat diet for 18 weeks. Each mouse had blood drawn at three times of day, each on a separate day: the first at Z10, the second at Z22, and the third at Z15. From these samples, we identified 907 lipid species from 16 lipid classes and subclasses, 223 polar metabolites, and 344 proteins. We found that diet, age, and blood-draw batch explained 44.3%, 15.6%, and 7.1% of the variability in the lipidomics data. In our metabolomics data, diet, age, and blood-draw batch explained 18.8%, 11.1%, and 8.1% of the variation, respectively. Following this analysis, we examined how each molecular species was associated with blood draw batch, age, and diet (Fig 3, Fig S5).

**Figure 3:**
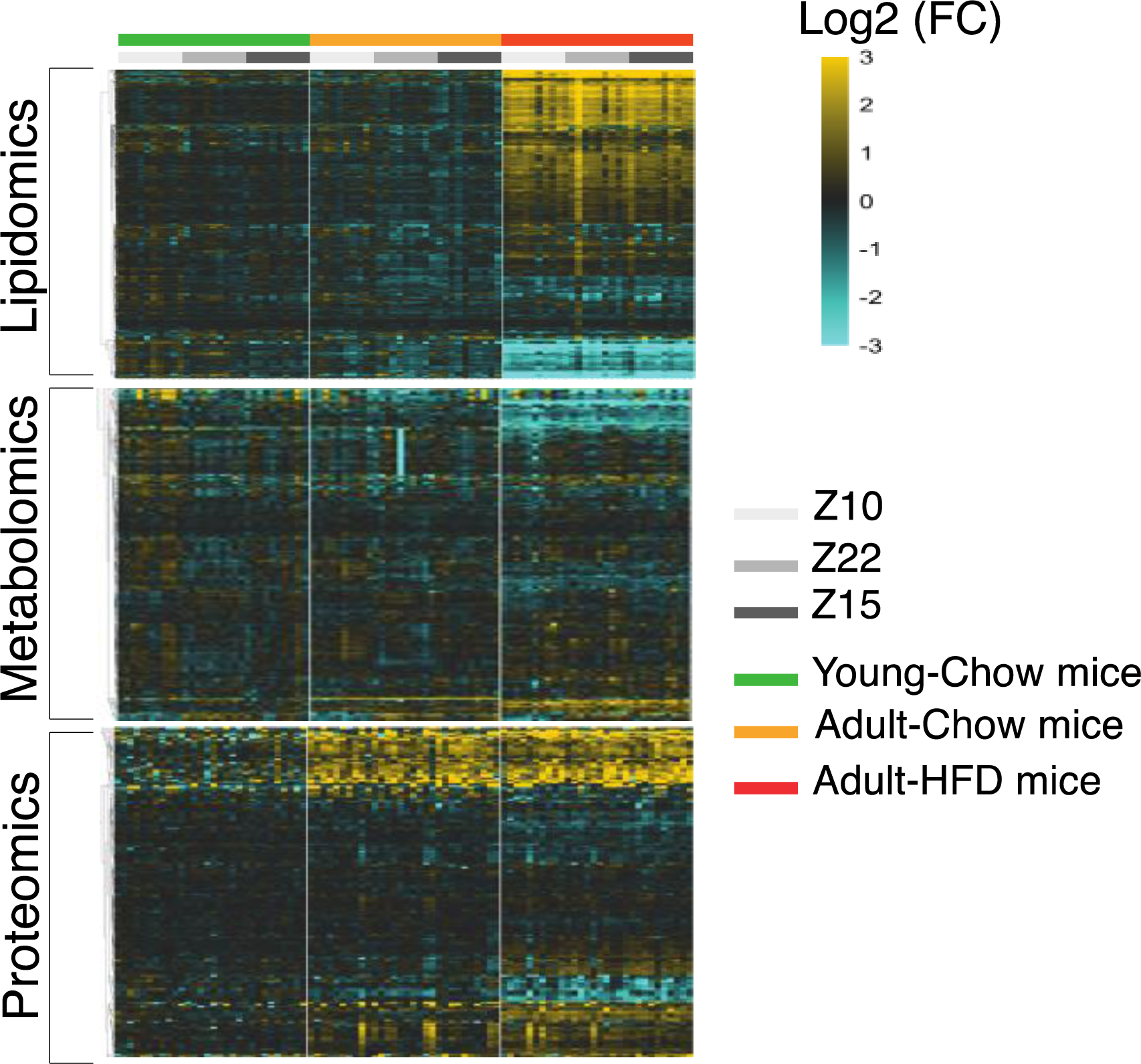
Heatmaps of metabolites, lipids, and proteins. Data was row-normalized to the mean value of the young chow-fed mice. Data displayed with hierarchical clustering by row for each data modality separately.

We noted that in chow fed animals at both ages, total TGs, glucose, and Apolipoproteins E (ApoE) and C-III (ApoC-III) were highest at Z10, while cortisol was lowest in the Z10 samples, all of which are previously reported to vary with food consumption (48–54) (Fig. 4). However, we also noted that these changes were muted, or absent, in the HFD-fed group, and the HFD group was the only one to show elevation of cholesterol esters in the Z22 and Z15 blood draw batches relative to Z10 (Fig. 4A).

**Figure 4:**
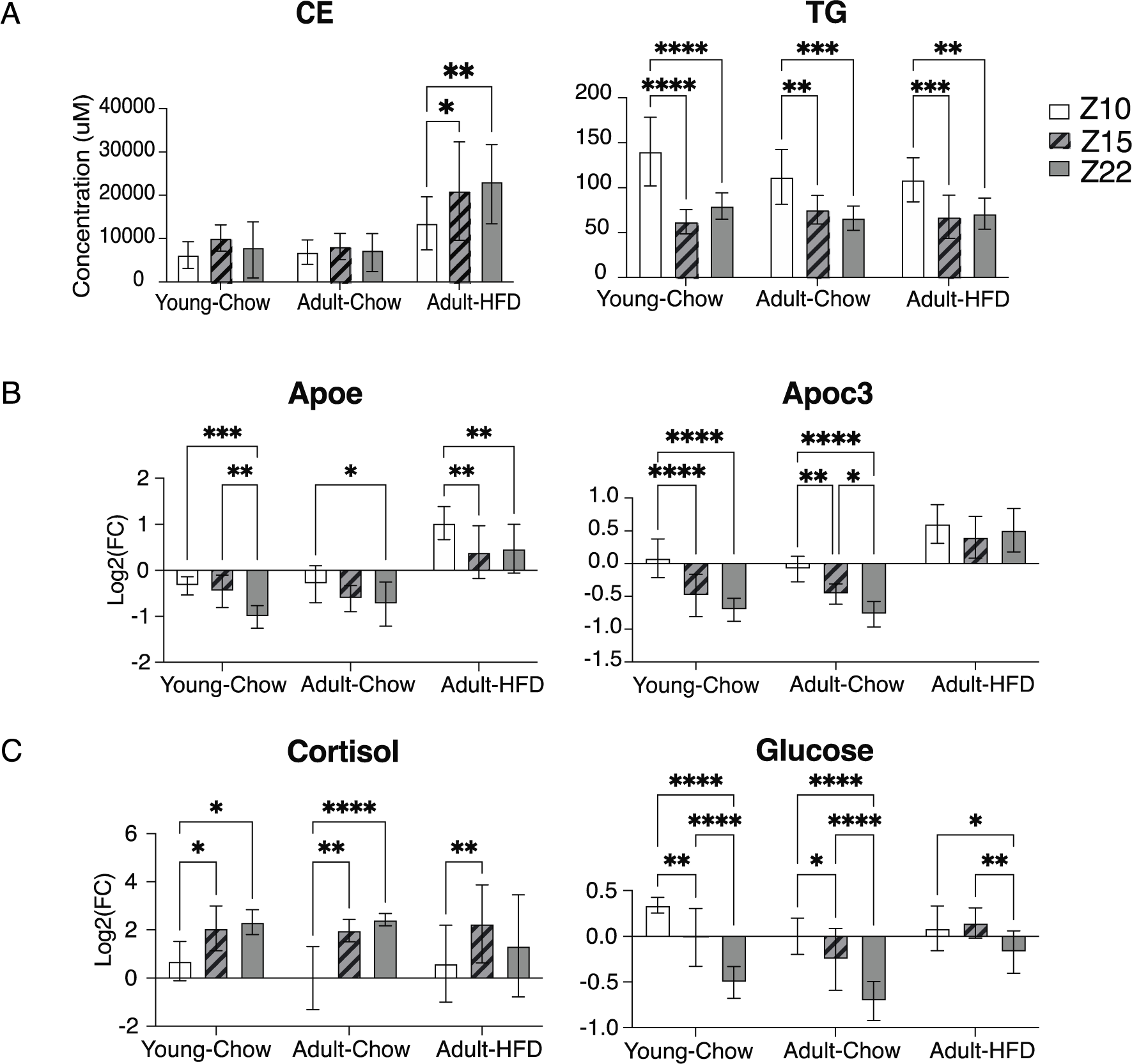
Selected changes observed between repeated blood draws with significance from two-way ANOVA analysis within each group of mice. (*** p <= 0.001, ** p <= 0.01, * p <= 0.05) A) Total concentration of TGs and CEs (n=10). Ammoniated adducts of TG and CE were normalized to their respective labeled standards from LipidSplash. The sum of concentration for all species per animal was calculated and presented here. Data shown is mean ± standard deviation. B) Relative concentrations of APOC3 and APOE at different blood draw time points (q<0.01). Protein concentrations were normalized to the bridge within each plex. Data shown is mean ± standard deviation. C) Glucose and cortisol levels change with time point in mice (q<0.01). The intensity of each sample was normalized to the mean signal of adult-chow mice at Z10 (n=10). Data shown is mean ± standard deviation.

#### 3.3.1 Age-associated changes in mouse plasma

We examined the molecular changes associated with age. We found that 39.1%, 12.5%, and 12.5% of lipids, metabolites, and proteins were significantly different between chow-fed animals at 8 and 24 weeks of age (q<0.01). As has been previously reported, we observed a small decline in the majority of lipid classes with age with LPCs, PCs, PIs, and TGs the most downregulated in our data. TGs containing 10-16 carbon acyl chains appear to be the most affected in that class, while other classes primarily showed lower levels of lipids containing FA 18:1, FA 22:6, FA 16:0, FA 14:0, especially the vinyl ether lipids p-16:0 and p-18:0 and choline headgroups (Fig. S6A).

We found 19 polar metabolites in the metabolomics data that decreased significantly with age (q<0.01), mainly N-acetylated amino acids, di-peptides (Glu-Glu, Glu-Ala), and modified amino acids. We found a modest decline of plasma glucose in the 24-week-old mice (55), with larger changes in ADP-ribose (>2-fold), and 4-hydroxycinnamic acid (> 10-fold) (56, 57). We identified 9 metabolites that were significantly increased, mostly amino acids, but also acetylcarnitine and the aging-implicated metabolite hippuric acid (58), which increased more than 10-fold between the 8-week and 24-week old chow fed groups (Fig. S6B).

In the proteomics data, we found that the immunoglobulins IgI, Ighm, IgG, Ighv were upregulated in adult mice (59), as were the 20S proteasomal subunit proteins Psma4, Psma5, Psma6, Psma7. We found a modest increase of Fibronectin (Fn1) in the adult mice, and a decrease in the pro-alpha1 chains of type I collagen (Col1a1), consistent with previous findings in human aged skin fibroblasts (58) (Fig. S6C).

#### 3.3.2 Diet-associated changes in circulating molecules

Finally, we examined differences in the plasma of the chow-fed and HFD-fed mice. Approximately 95% of lipid species (863 of 907) were significantly different between the two diets (q < 0.01), and there was an increase in total lipid content, consistent with previous reports (60, 61). The most profound change was in cholesterol esters, which accounted for 96.6% of the total molar concentration difference, with one particular species, arachidonoyl-cholesterol (CE 20:4), accounting for approximately 65% of the total molar increase in lipids in HFD-fed mouse plasma. Non-intuitively, but consistent with previous reports, we found that the high-fat diet did not increase the total circulating triglyceride concentration (62, 63).

Lipid class enrichment analysis showed that SMs, PCs, and CEs were the most highly increased lipid classes in the high-fat diet plasma, while PIs, TGs and DGs were the most depleted (Figure S7A). Enrichment analysis on fatty acyl composition found that polyunsaturated fatty acids were highly perturbed, with an increase in most ω-6 fatty acyls and a decrease in most ω-3 fatty acyls (Figure S7B).

The total ω-6/ω-3 ratio increased from 1.18 ± 0.05 in adult chow mice to 2.40 ± 0.1 in HFD mice, consistent with previous studies on obesity and type II diabetes in mice (64, 65). Although this data does not resolve double bond positional isomers, we annotated ω-6 and ω-3 lipids were based on the most abundant isomer form in mouse plasma, as previously reported via Fatty Acid Methyl Esters (FAME) analysis of mouse plasma and serum (66, 67).

Next, we examined the total molar concentration for each acyl chain in each lipid class. Lipids containing arachidonic acid (AA, FA 20:4, ω-6) and eicosanoic acid (EPA, FA 20:5, ω-3) were found in most classes and were consistently changed by diet, with AA significantly higher in every lipid class and EPA lower in all lipid classes in which they were found (Fig.5). EPA was the only ω-3 lipid that decreased globally, while its main product, the more abundant decosahexanoic acid (DHA, FA 22:6, ω-3), increased in all lipids classes except PIs, DGs and TGs. The ω-6 fatty acid AA and its product adrenic acid (FA 22:4, ω-6) increased significantly in every lipid class in which it was detected.

**Figure 5:**
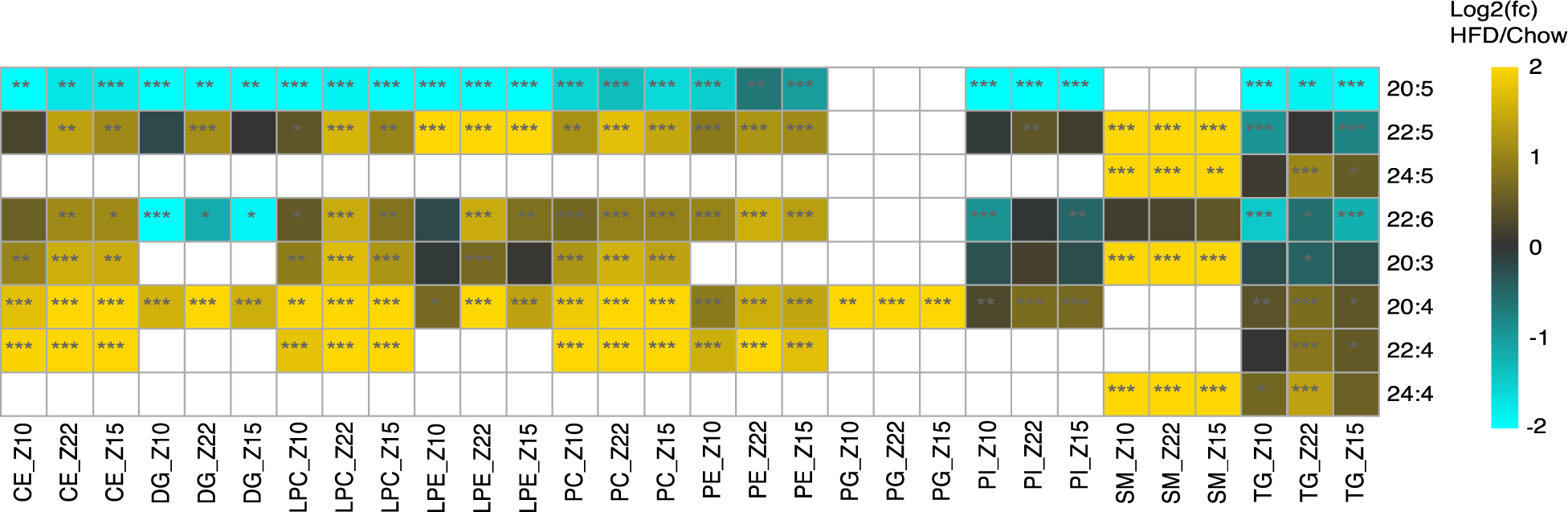
Heatmap of the average of log_2_ (HFD/ Chow) total concentration of PUFA acyl chains for every quantified lipid class at each timepoint. p-values were calculated using a two-tailed Student’s t-test to differentiate adult-HFD-fed mice from adult-chow-fed mice. (*** p <= 0.001, ** p <= 0.01, * p <= 0.05). Missing values are shown as white boxes.

In the metabolomics data, we found that 54.3% of measured metabolites were significantly different between the HFD- and chow-fed mice (q<0.01). Metabolomics data demonstrated that the free fatty acids, AA and EPA, were changed in the same direction as the complex lipids containing them did. Free AA increased more than 4-fold, and EPA dropped more than 6-fold. We also found a slight increase in the glycerophospholipid-related metabolites glycerol-3-phosphate and glycerophosphocholine, while choline, acetylcarnitine, and carnitine levels consistently declined in the high fat diet (Figure 6A).

**Figure 6:**
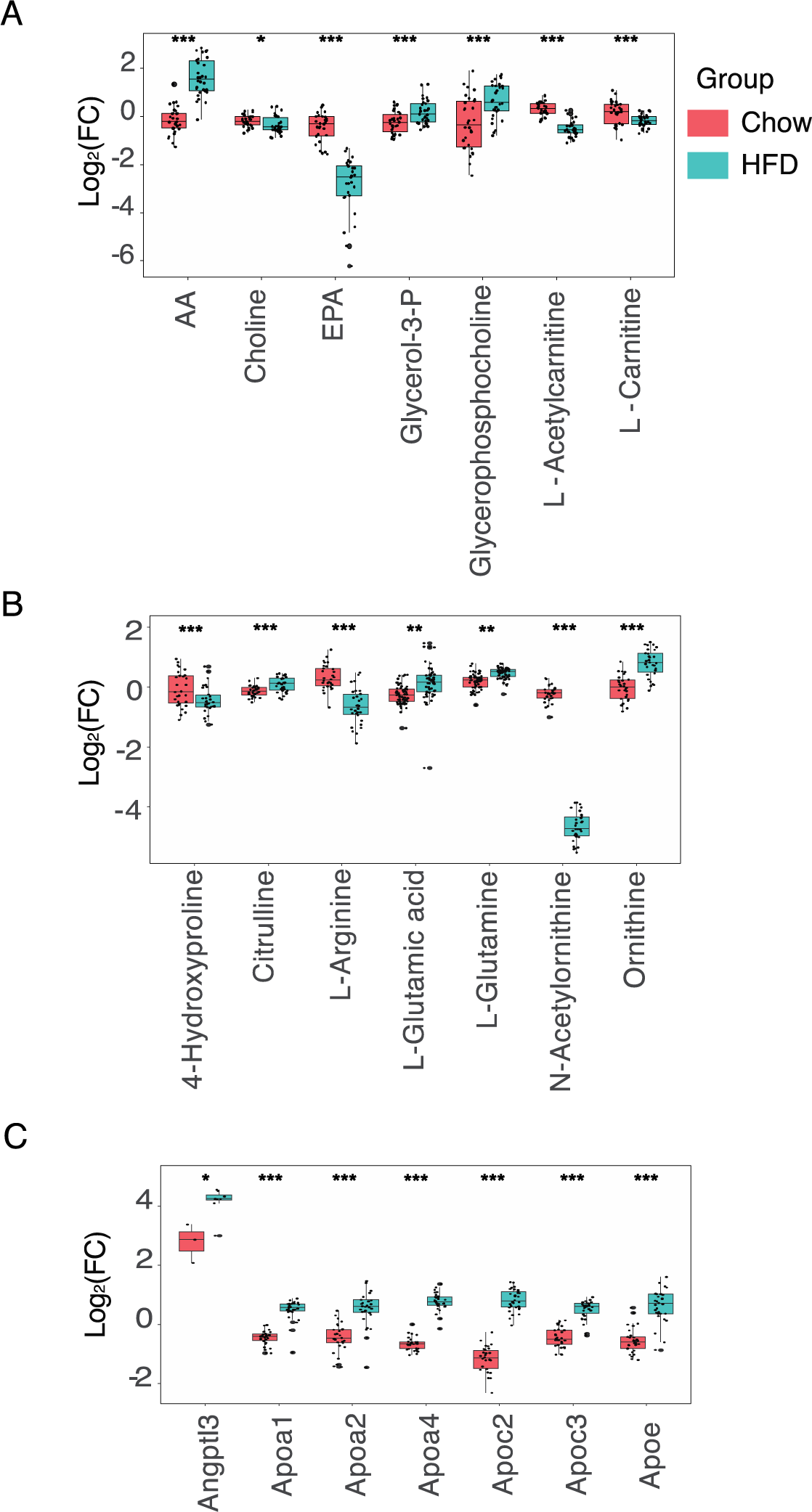
Significant differences in plasma metabolites and proteins of high fat diet fed mice. A) Polar metabolites associated with lipid metabolism showed a dietary effect (*** p-value <0.001, * p-value <0.05). B) Significant metabolites in nitrogen metabolism changes with diet (*** p-value <0.001, ** p-value <0.01) C) Lipoproteins were upregulated in HFD-fed mice (q<0.01). Proteins were normalized to the bridge included in every plex.

We noticed that many nitrogen metabolism-related metabolites were significantly different in HFD-adult mice (Fig.5B). In particular, citrulline, glutamate, glutamine, and ornithine were increased, while 4-hydroxyproline, arginine, and N-acetylornithine were decreased, suggesting possible impairment of the urea cycle or high activity of hepatic Arginase I, which is reported to be induced in obesity (68).

Finally, we found that 21.5% of measured proteins significantly differed between diets (q<0.01). We observed significant upregulation of many apolipoproteins (Angptl3, Apoa1, Apoa2, Apoa4, Apoc2, Apoc3, Apoe) (Fig.6C), the H-2 class I histocompatibility complex (H2-Q10/Q4), and cytosolic malic enzyme (ME1), all in agreement with previous findings (69–73). We also found that the previously identified markers of obesity fructose-biphosphate aldolase B (ALDOB), betaine-homocysteine methyltransferase (BHMT), and haptoglobin (Hp) were upregulated more than 4-fold in the HFD mice (74), and that several members of the complement system (Cfd, C8g, C8a, C8b) were significantly depleted.

## 4. Discussion

### 4.1 Development of automated LLE: evaluations of linearity and reproducibility

Here we present, for the first time, an automated sample preparation system using an MTBE-based liquid-liquid extraction on a PAL-DHF followed by Auto-MP3 protein preparation for high quality multi-omic analysis of metabolites, lipids, and proteins from a single blood plasma sample, enabling a deep phenotyping and an increased understanding of system biology. This method offers a robust workflow to prepare samples for multi-omic analysis via mass spectrometry with minimal human intervention.

We selected the MTBE-based extraction approach, based on the method of Matyash et al., which demonstrated excellent recovery of lipids while avoiding chlorinated solvents, and resulted in the automation-friendly location of the protein pellet at the bottom of the extraction vessel. We focused on setting up a system in which the consumables are available off-the-shelf from major laboratory suppliers to make implementation easy for the community while providing high-quality data. We created an automated workflow that avoids the issues of contamination and sample loss that can occur when lipids and organic solvents are exposed to plasticware, which can cause ion suppression and reduce numbers of lipid identifications as mass spectrometer scans (43, 44).

We demonstrated the accuracy and precision of this platform for lipids, which was comparable to other publications performing an LLE-based lipid extraction (45, 46). We also found comparable accuracy and precision for metabolites from the aqueous phase, a much less studied group of compounds from this workflow (20, 75, 76). We had a higher CV in the measurement of citrate, a tri-carboxylated metabolite that suffers from substantial peak broadening during chromatographic separation. We suspect this is a chromatographic artifact, as it is difficult to optimize chromatography for the diverse classes of compounds in metabolomics. The determined ratio of labeled:unlabeled standards showed a highly accurate recovery of lipids and metabolites. We found that this method was most reproducible on phospholipids, while the more hydrophobic lipids (DG, FA, MG, PG, and TG) had higher, but acceptable variability. Previous literature found that recovery of non-polar plasma lipids using MTBE extraction protocol was lower than for phospholipids (45, 47), suggesting this may be unavoidable when performing MTBE-based LLE. These neural lipids are also extracted poorly in one-phase extractions when using a higher content of polar solvent (77). If neutral lipids, especially TGs are of paramount importance to an experiment, other protocols have been shown to have higher recoveries of TGs, including the 3PLE hexane phase (78) or hexane-isopropanol extraction (79) or ethyl acetate-ethanol (80), but at the expense of reduced phospholipids recovery in the same phase.

In addition to its high accuracy and reproducibility, this methodology offers substantial time savings and requires only a small sample volume to prepare metabolomics, lipidomics and proteomics samples. The method we have implemented on the PAL-DHR can extract 90 plasma samples in less than 24 hours, preventing it from being a bottleneck if mass spectrometry data acquisition takes greater than 15 minutes per sample, while recovering a similar number of identifications to manual preparation. In contrast, careful manual extraction without exposure to plasticware for 90 samples would take three 8-hour days.

This is the first report of an integrated system that uses only glassware and stainless steel consumables with a centrifuge and cooling system that can automate every step of the liquid-liquid extraction (LLE) while minimizing contaminations, to our knowledge. We have carefully described the system, the pitfalls we encountered in implementing automated sample preparation with it, and optimized consumables to enable other laboratories to create a similar workflow.

### 4.2 Changes in the plasma lipidome, metabolome, and proteome induced by a high-fat diet

Using the PAL-DHR prepared samples, we successfully recovered a number of previously reported biomarkers associated with age and high-fat diet (HFD) in mice, demonstrating the ability to recover biological signals from real samples. Additionally, the fact that we could recover the dramatically different lipid content in the plasma of HFD-fed mice demonstrates that the method is robust to samples with strikingly different lipid concentrations, while also capable of finding subtle changes, like those between blood draw time points.

Although we observed some molecular concentration differences between blood draw batches that are consistent with known circadian effects (such as glucose and total triglycerides), an important caveat to the data is that the different time points were sampled in separate weeks, and some of the detected changes could be due to week-to-week variation as opposed to circadian changes, limiting the interpretability of novel findings around circadian rhythm from this data set. Hence we chose to focus on examining effects of circadian changes that had been previously reported.

As expected, given the significant physiological changes induced by a high-fat diet (62, 81–84) we found that the differences in molecular profiles between diets was much larger than all other effects in the data set. Lipids were the most perturbed class of molecules, consistent with the increased dietary lipid content. Proteomics results showed elevated levels of many different apolipoproteins (APOA4, APOC2, APOE, APOA1, APOA2, and APOB), important to the circulation of the increased lipids load. We observed changes in the polar compounds that are both the precursors and products of lipid head groups, suggesting that there may be changes in the metabolic fluxes in lipid metabolism in addition to the differing levels of circulating lipids. As expected, we found elevated glucose levels in the HFD mice, consistent with insulin resistance comitant with diet induced obesity. We found lower acetoacetate levels and a shift in the circulating pyruvate/lactate ratio, suggesting that the intracellular NAD/NADH ratio may be reduced by the high-fat diet (85). Significantly, we noticed a reduction of PI lipids containing docosahexaenoic acid (DHA). PIs are known to be the main donor of AA to produce downstream bioactive lipid mediators (86, 87), but DHA-containing PI are less characterized but could be interpreted as depletion in the precursor pool of DHA for the production of pro-resolving mediators in a high-fat diet.

Pakiet et al. previously reported that a high-fat diet increased the levels of total AA and DHA acyl groups as determined by GC-MS based analysis of Fatty Acid Methyl Esters (FAME) (61). Our analysis showed that CEs are at a substantially higher concentration than other circulating lipid classes (excluding free fatty acids, which were not quantified in this study), consistent with previous reports (62). Hence, FAME measurements of the fatty acid composition of plasma is primarily an analysis of the acyl groups from CEs. We report here for the first time arachidonic acid is increased and eicosapentaenoic acid is depleted uniformly across all major plasma lipid classes, despite the fact that the vast majority of the change in the total AA:EPA ratio is from arachidonoyl-cholesterol (CE 20:4). We also demonstrated that the major change in ω-3 fatty acids is the depletion of EPA, while DHA was upregulated in some classes, and downregulated in others. Consistent with EPA depletion having an important role in the negative effects of a high-fat diet, it has been previously reported that animals fed a high-fat diet supplemented with EPA have improved glucose tolerance, reduced liver adiposity, and decreased weight gain when compared to an unsupplemented high-fat diet (88) (92). This acyl-composition shift to increased AA and depletion of EPA is present in phospholipids as well, providing a larger source of AA for eicosanoid synthesis. In the metabolomics study, we confirmed that there was a relative increase in AA, and depletion of EPA free fatty acids in HFD mice.

We hope that this comprehensive analysis of effects of age and diet on the plasma multiome will serve as a resource for the community working on these topics.

## 5. Data availability statement

Metabolomics and Lipidomics data will be available from Metabolomics Workbench: DataTrack ID: 4147

Proteomics raw files will be available from the PRIDE database PRIDE PXD049202

## Acknowledgments

The authors would like to thank Tom Tobien from Trajan for the initial setup of the instrument, and Dr. David Botstein for helpful comments on the paper while it was being drafted. We also acknowledge the contributions of Dr. Niclas Olsson for acquiring the proteomic samples, and Travis Lee for assembling the enclosure for the robot.

## Supplement Figures

**Figure S1:**
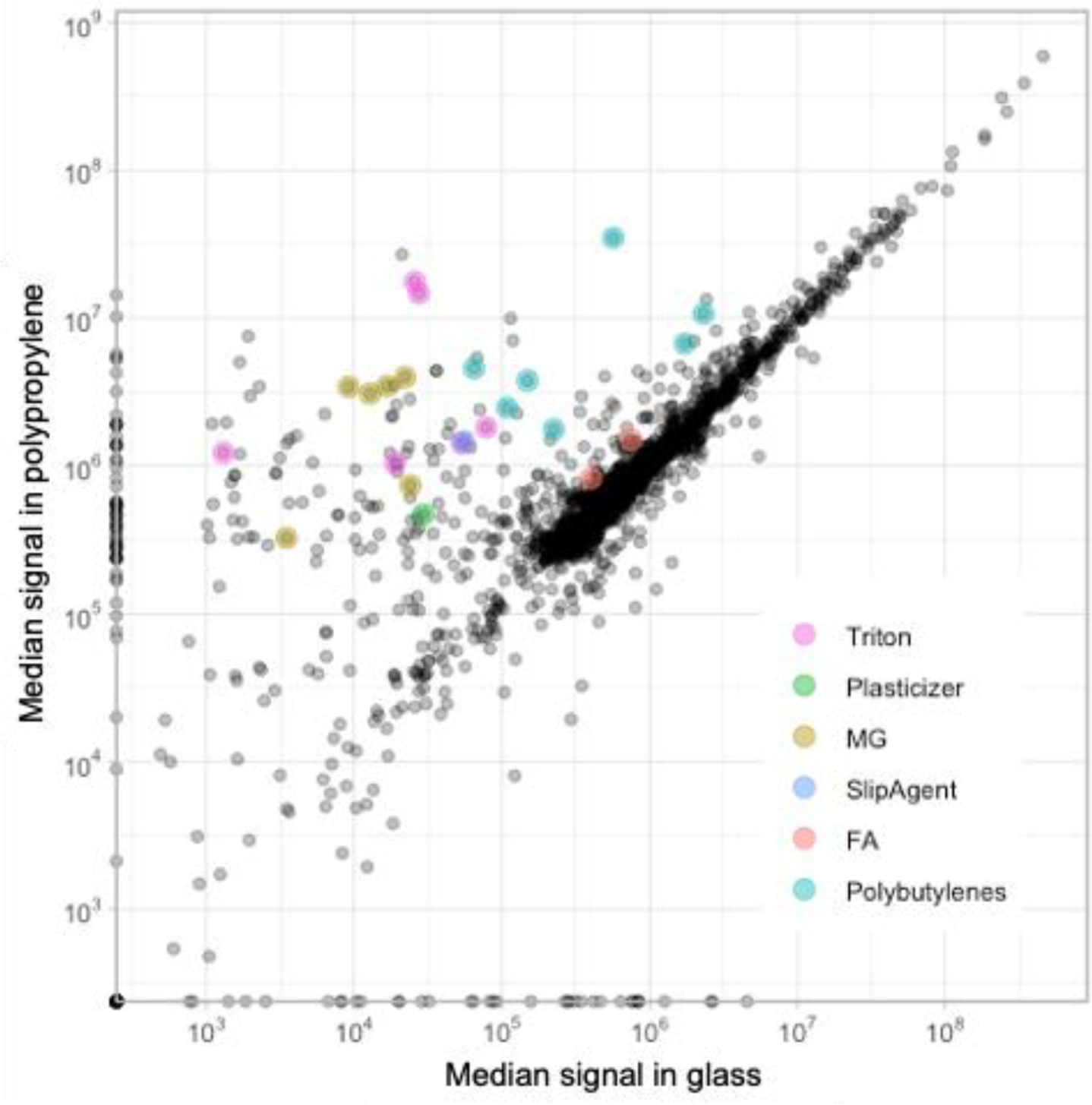
Scatter plot of mass spectrometry features from samples manually extracted in glass vials vs. polypropylene tubes (n=3).

**Figure S2:**
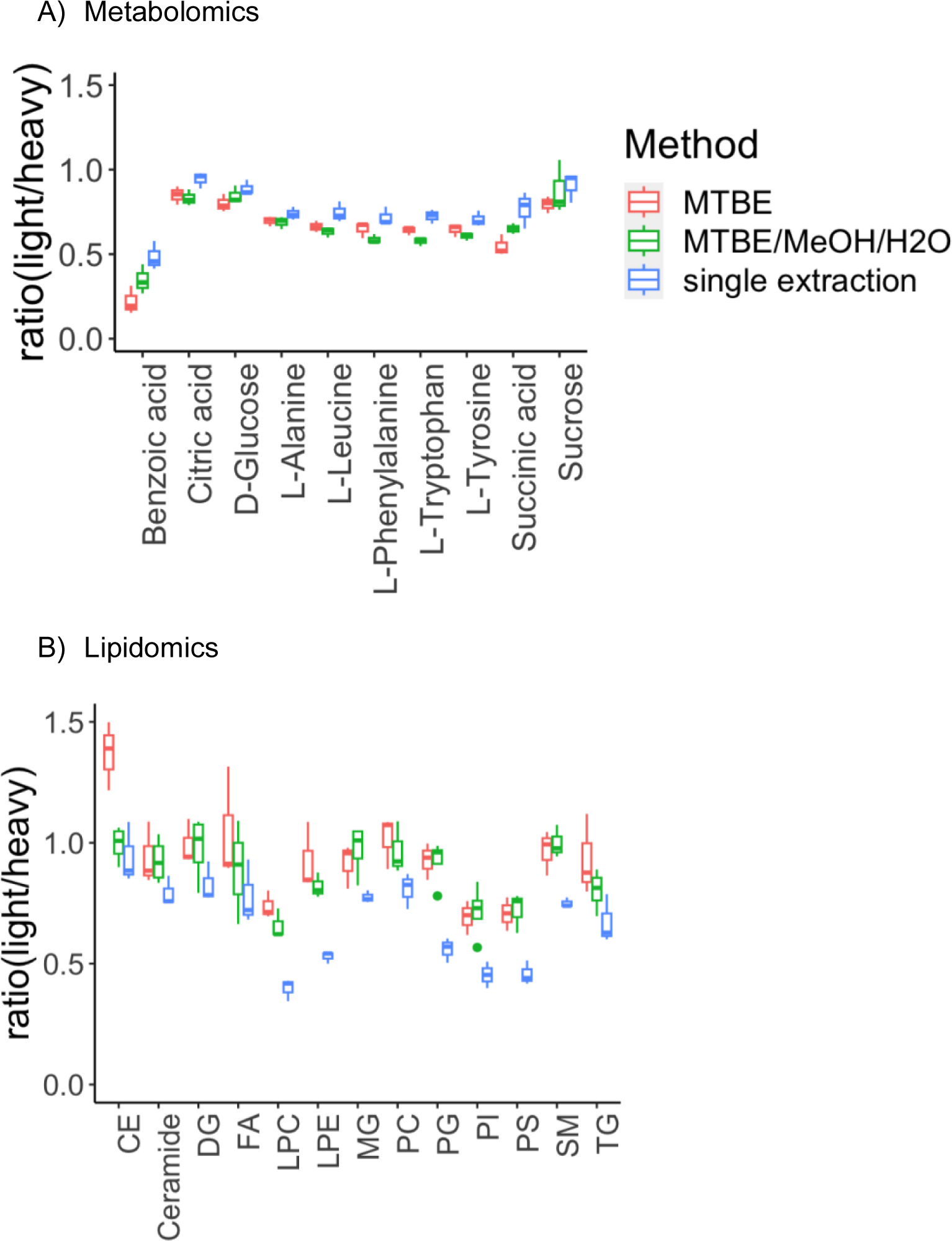
Recover efficiency of lipids and metabolites after LLE (n=3). Red= 100% MTBE, Green= MTBE/MeOH/H_2_O (3.33:1:1,v/v/v), Blue = without second extraction.

**Figure S3A:**
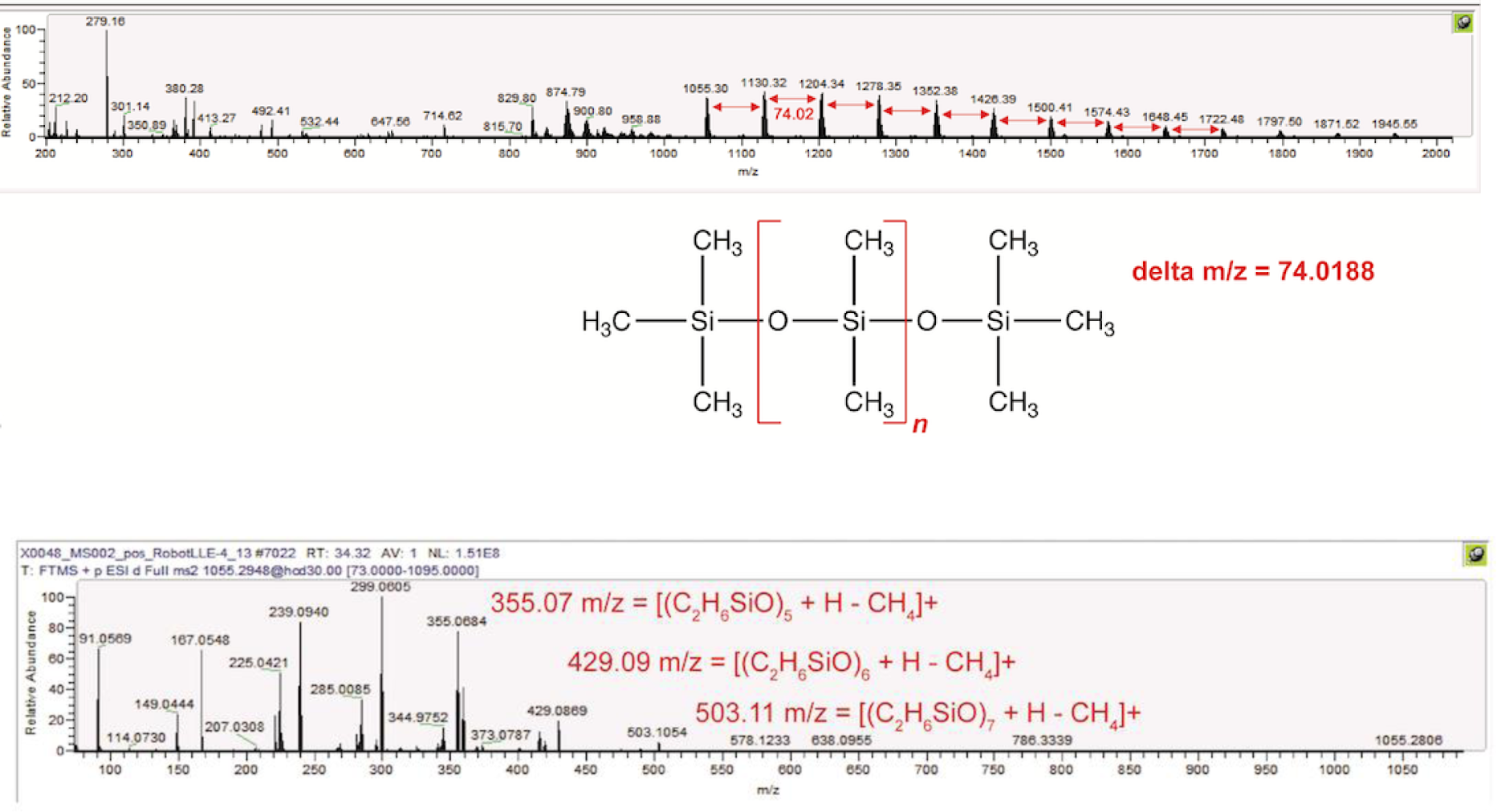
Polysiloxane EIC and MS/MS when using blue silicone cap - Top-MS1, bottom-MS2. Identification of polysiloxanes was done by matching their exact masses and the 74.02 D spacing between peaks in both the MS^1^, and the MS^2^ fragments of each precursor.

**Figure S3B:**
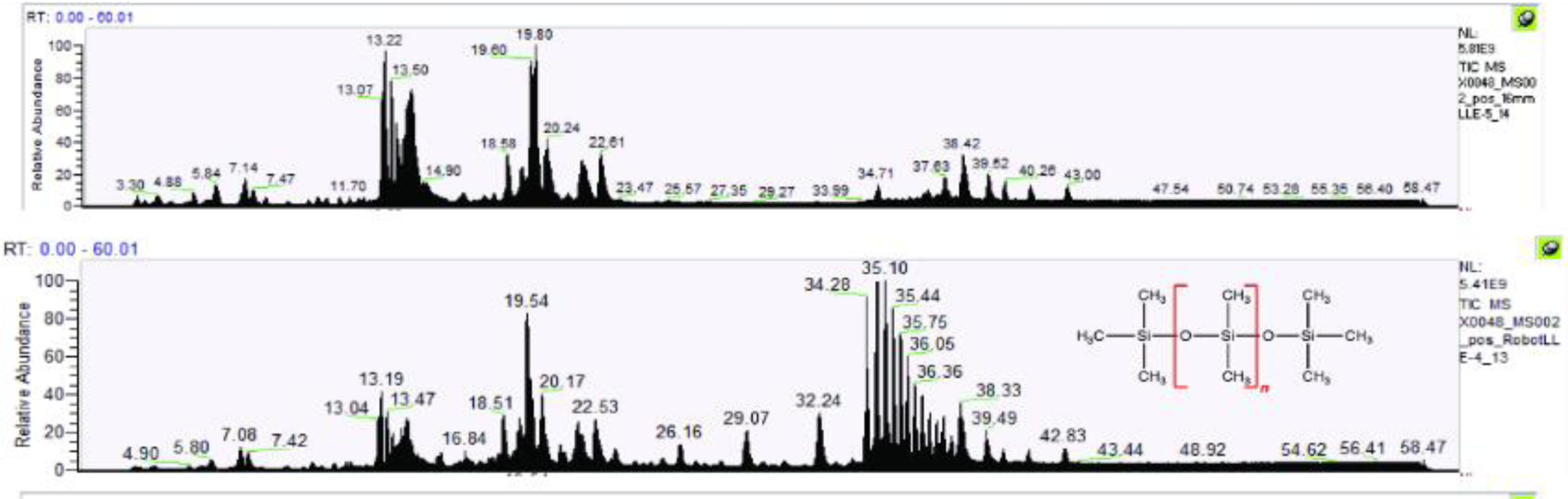
MS1 spectra of lipid data with PTFE disc (Pure_PTFE) (top) and silicon cap (bottom)

**Figure S4A:**
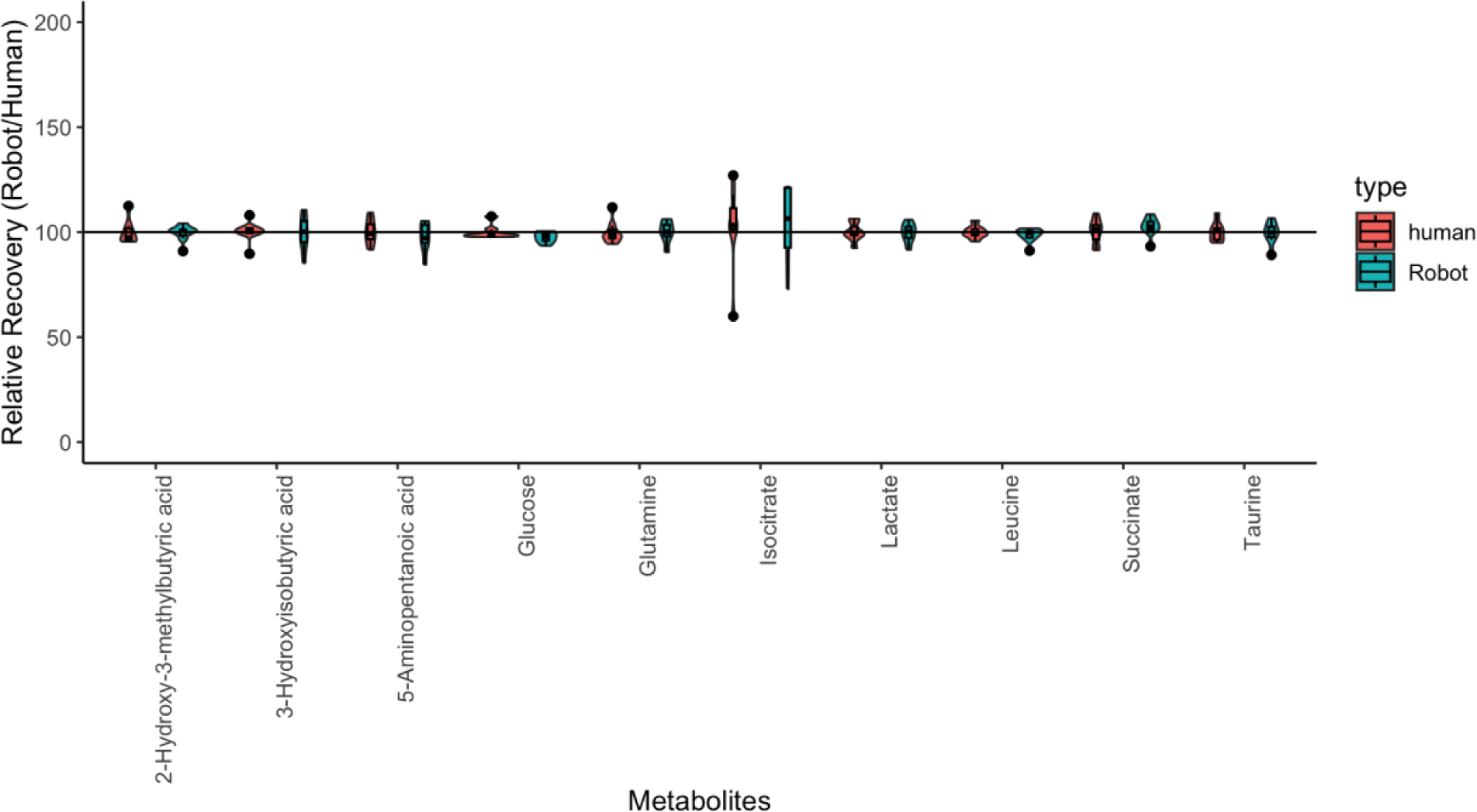
Relative recovery of the 10 plasma metabolites with the highest intensity. 10μL of mouse plasma was extracted manually (n=10) or by robot PAL (n=7). PeakAreaTop of each plasma metabolite extracted by the robot was normalized to the mean of PeakAreaTop of the manual extractions.

**Figure S4B:**
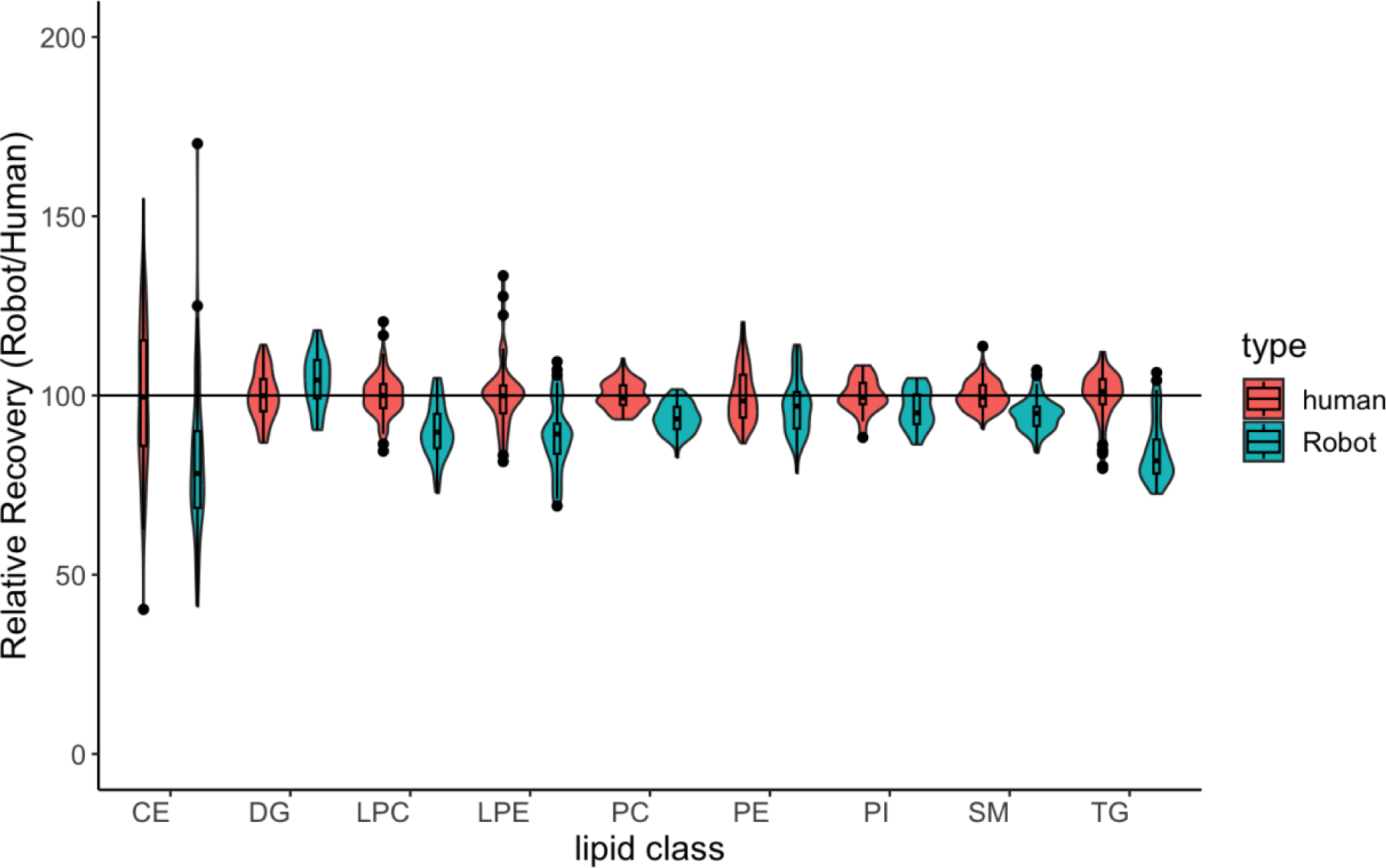
Percent recovery of Top 10 plasma lipids per lipid class. 10μL of mouse plasma was extracted manually (n=10) or by robot PAL (n=7). LipidSplash standards were spiked in post-LLE, during resuspension. Peaks were normalized to the class-specific LipidSplash standard. Relative recovery represents the concentration of lipids relative to the mean of the manually extracted lipids from that class.

**Figure S4C:**
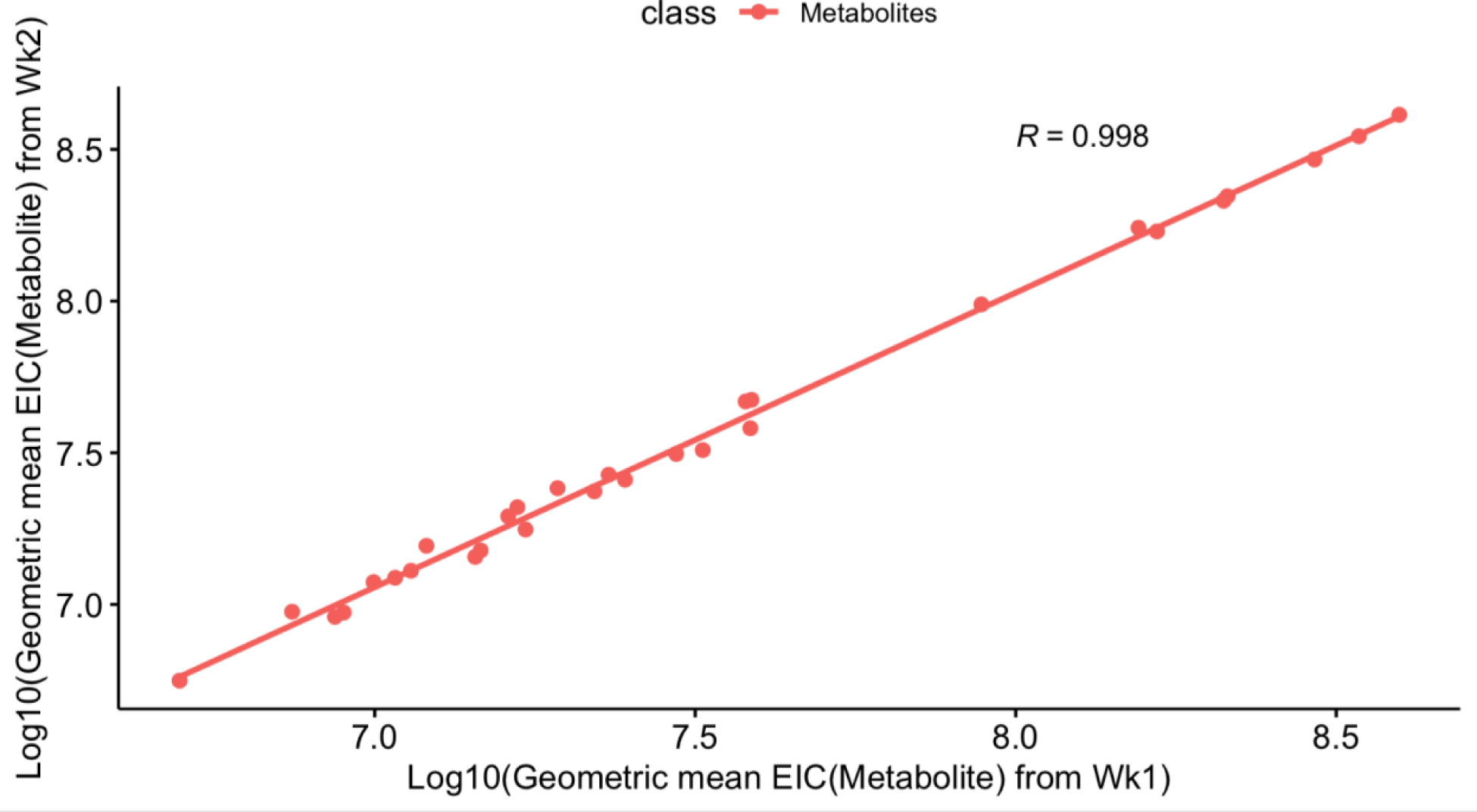
The correlation between the top 10 most abundant plasma metabolites (20, 30, and 50 μL plasma, n = 3) is represented in a scatter plot. The x-axis denotes the natural logarithm of the geometric mean of peak area obtained in week 1, while the y-axis represents the natural logarithm of the geometric mean of peak area collected in week 2. Each dot on the plot corresponds to the mean log-transformed peak area of a specific metabolite per unit of plasma volume.

**Figure S4D:**
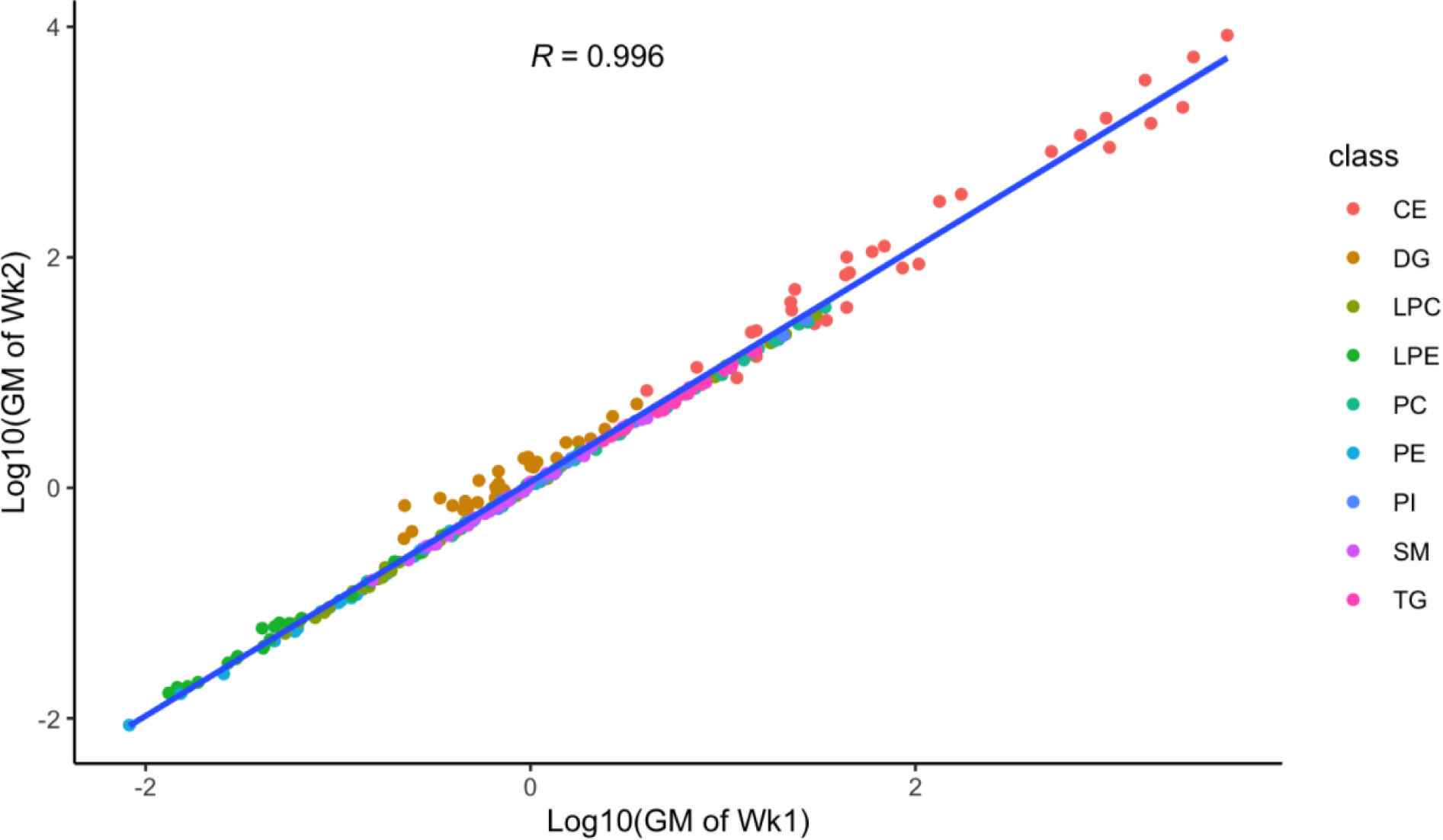
Correlation between the top 10 most abundant plasma lipids from each lipid class, as extracted using automated liquid-liquid extraction (LLE) from three samples of plasma volumes (20, 30, and 50 μL). The peak areas of each analyte were normalized to the peak area of the corresponding LipidSplash standard, followed by natural logarithm transformation. The x-axis represents the natural logarithm of the geometric mean of lipid concentration (nM) obtained in week 1, and the y-axis represents the natural logarithm of the geometric mean of lipid concentration (nM) obtained in week 2. Each dot represents the average log-transformed quantification of unique lipids per plasma volume. The lipid classes are represented by different colors.

**Figure S4E:**
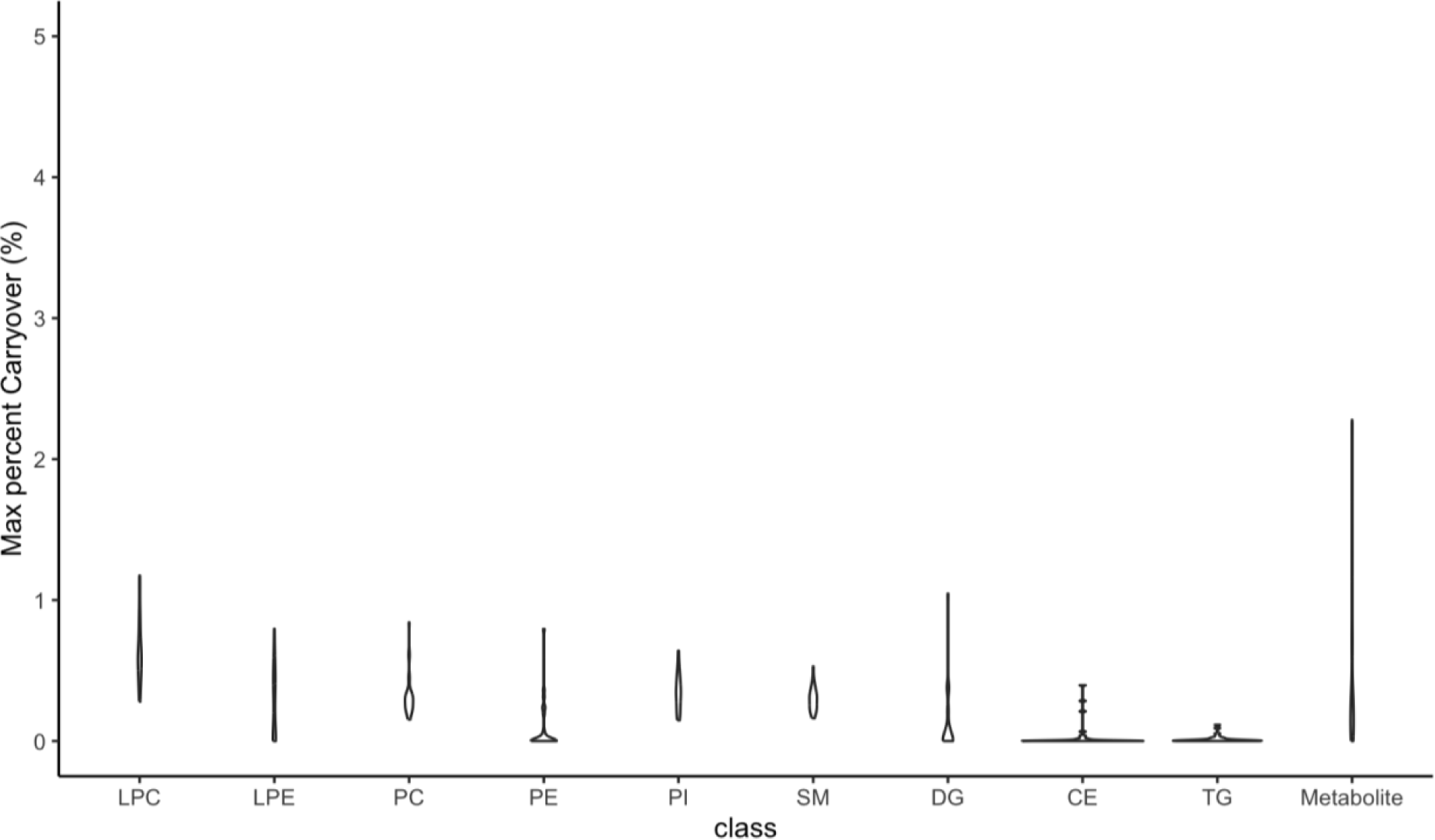
Percent carry over of top 10 most abundant plasma metabolites and lipids per lipid classes. Samples were arranged and extracted with a blank in between each plasma sample (n=3) at various plasma volumes (20, 30 and 50 μL). The carry over for metabolites and lipids was calculated as the fraction of the peak area of the analyte in the blank compared to the plasma injected before it.

**Figure S5A:**
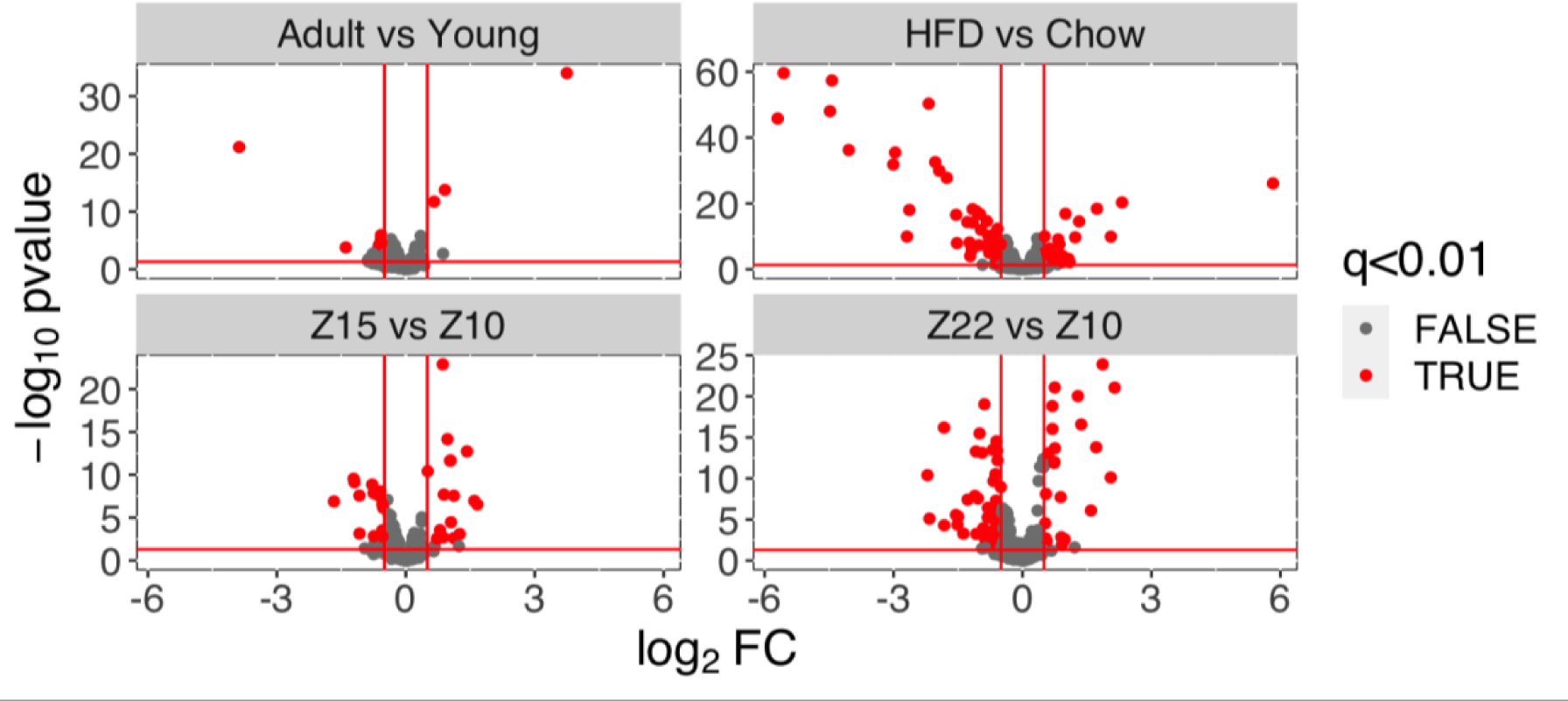
Significant metabolites changed with time point, age, and diet (q<0.01)

**Figure S5B:**
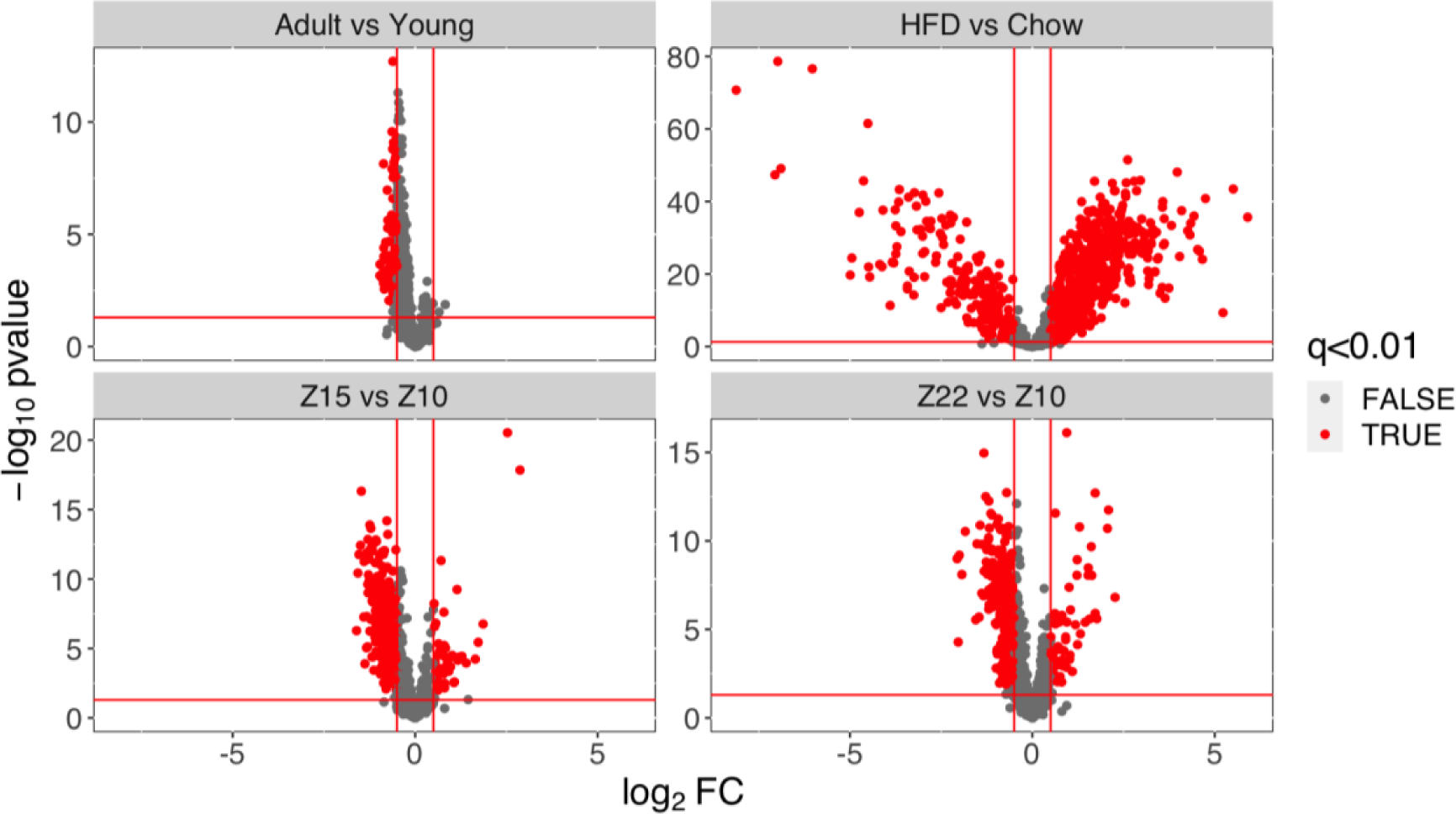
Significant lipids changed with time point, age, and diet (q<0.01)

**Figure S5C:**
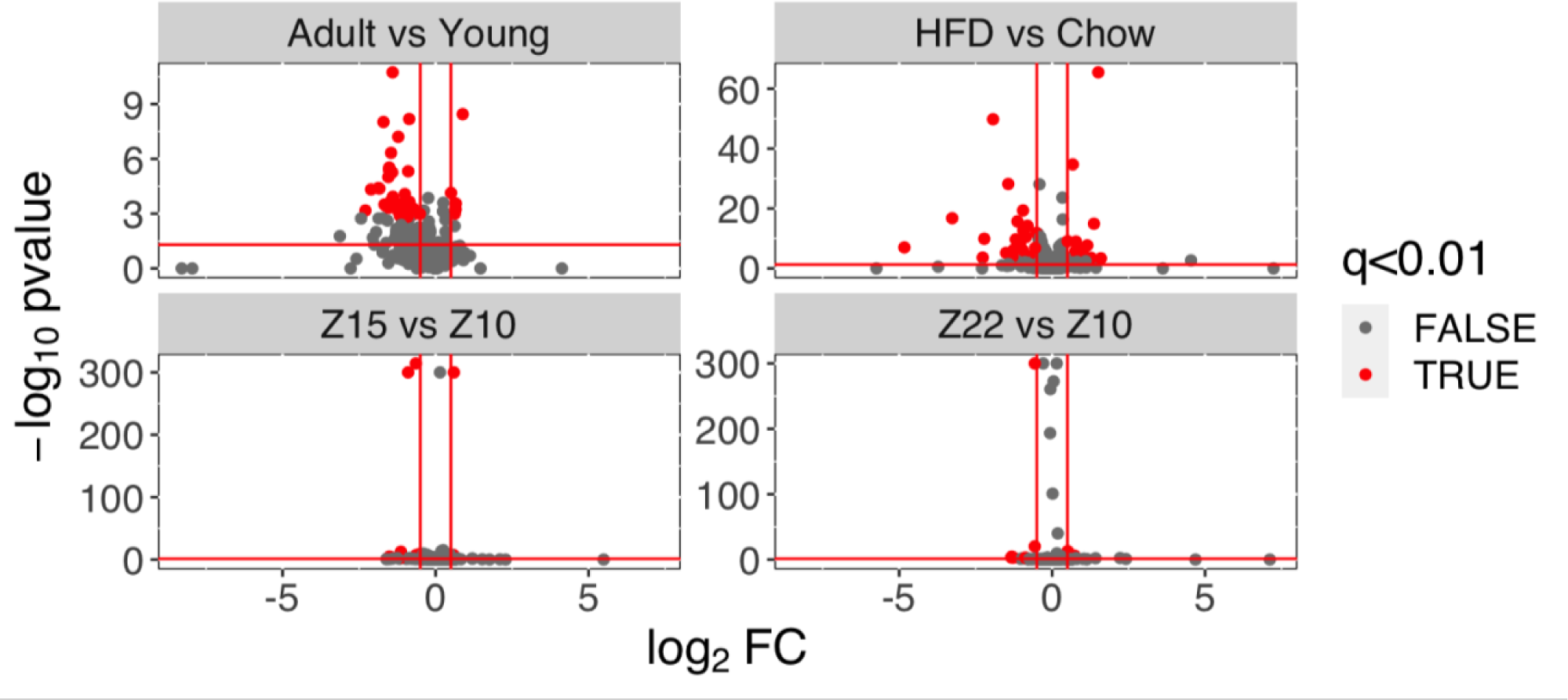
Significant proteins changed with time point, age and diet (q<0.01)

**Fig.S6A:**
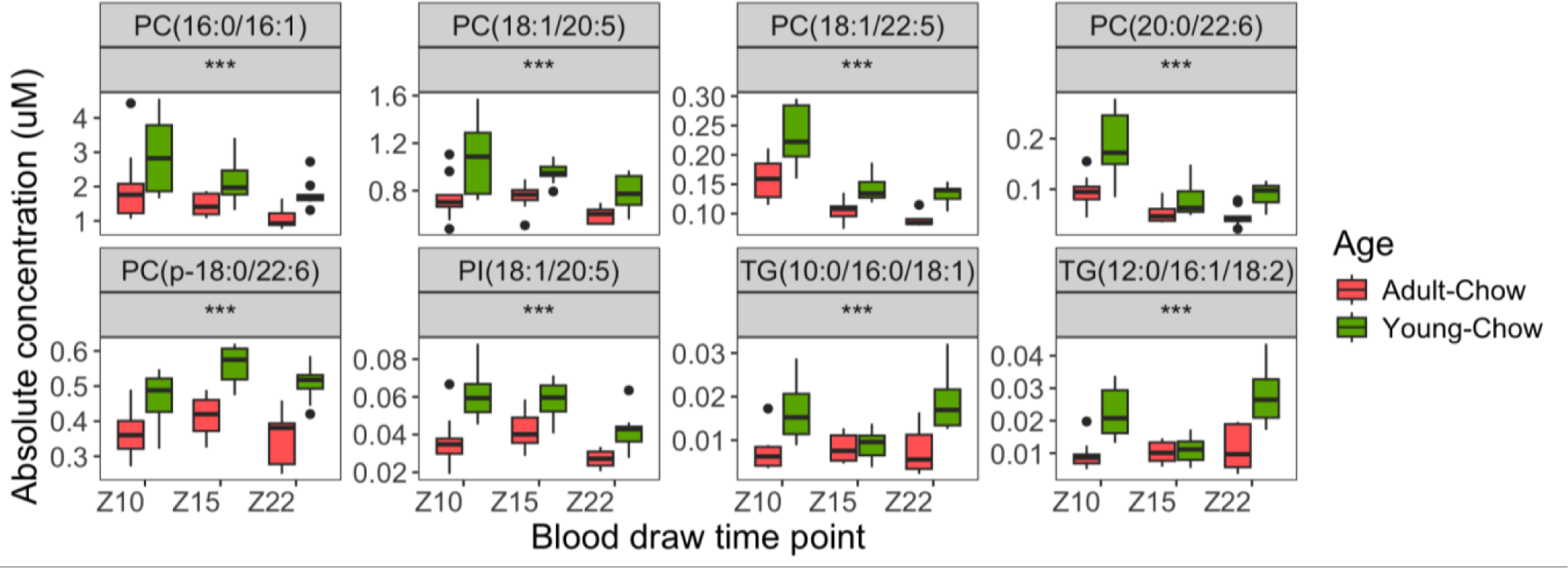
Selected lipids showing a significant age effect (*** p-value <0.001, * p-value <0.05) between the young-chow (green boxes) and adult-chow (red boxes) groups.

**Fig.S6B:**
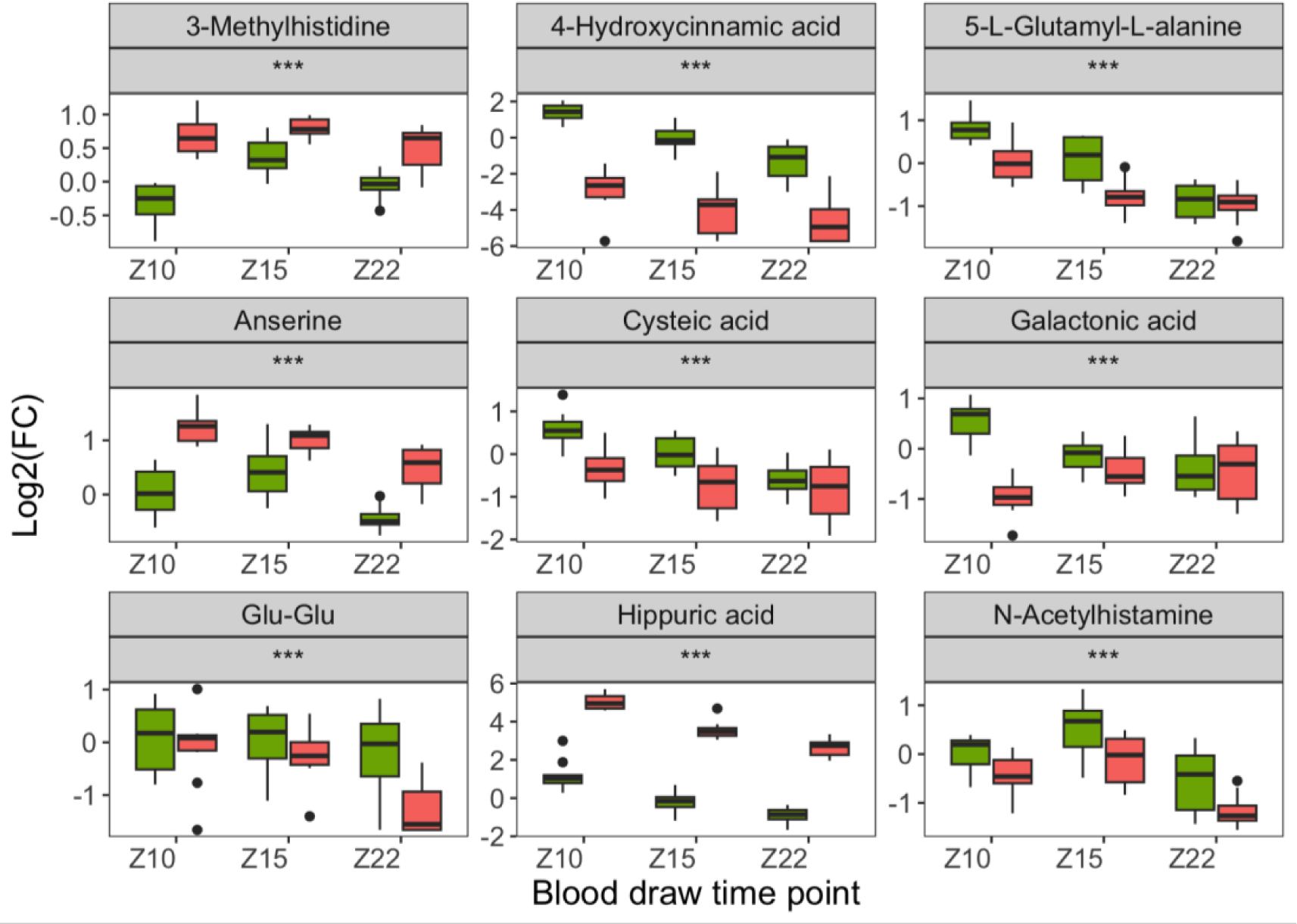
Selected metabolites showing a significant age effect (*** p-value <0.001, * p-value <0.05) between the young-chow (green boxes) and adult-chow (red boxes) groups. Value are normalized the the mean value in adult-chow at Z10.

**Fig.S6C:**
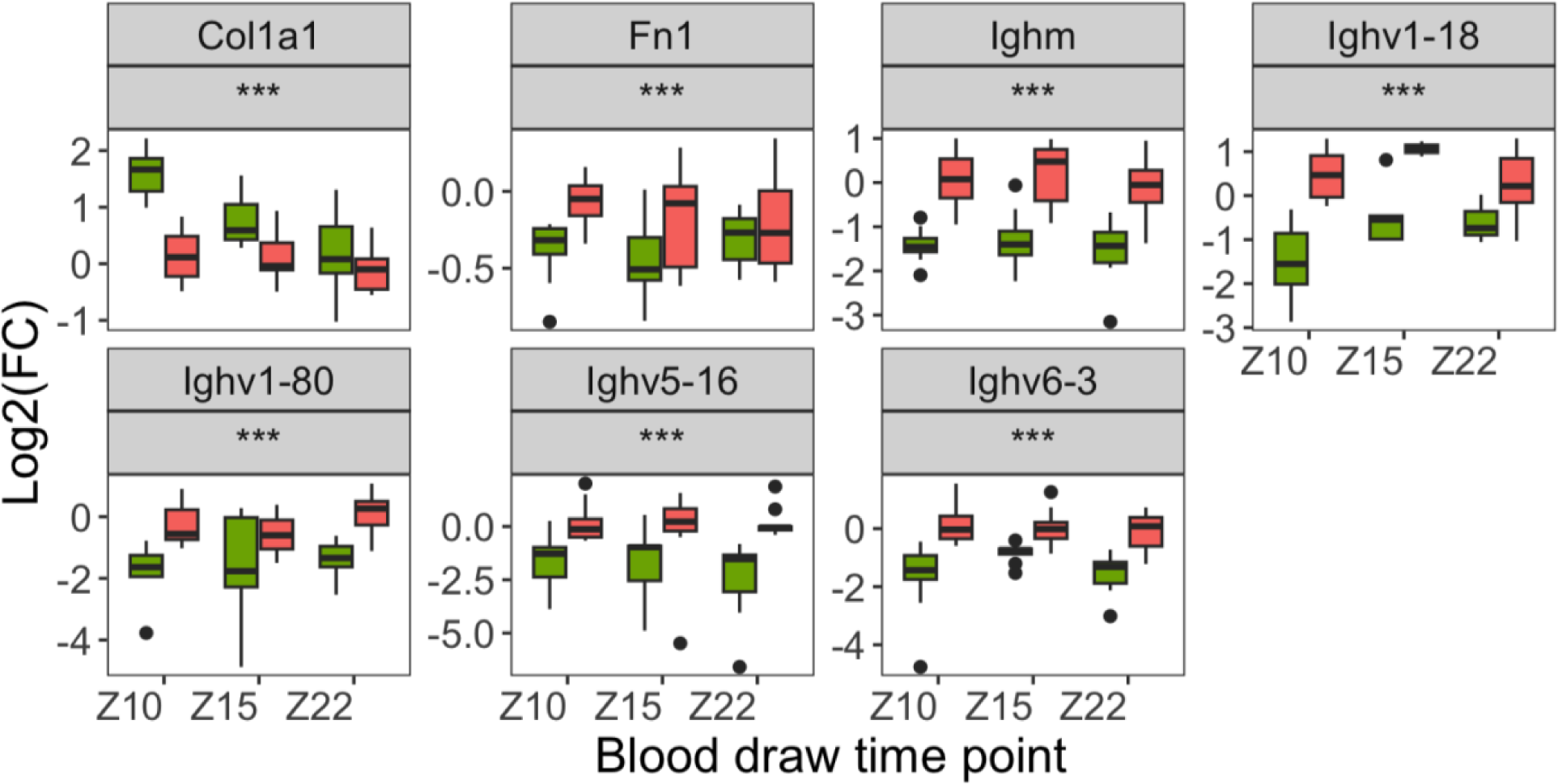
Selected proteins showing a significant age effect age effect (q<0.01,*** p-value <0.001, * p-value <0.05) between the young-chow (green boxes) and adult-chow (red boxes) groups. Proteins were normalized to the bridge included in every plex.

**Figure S7A:**
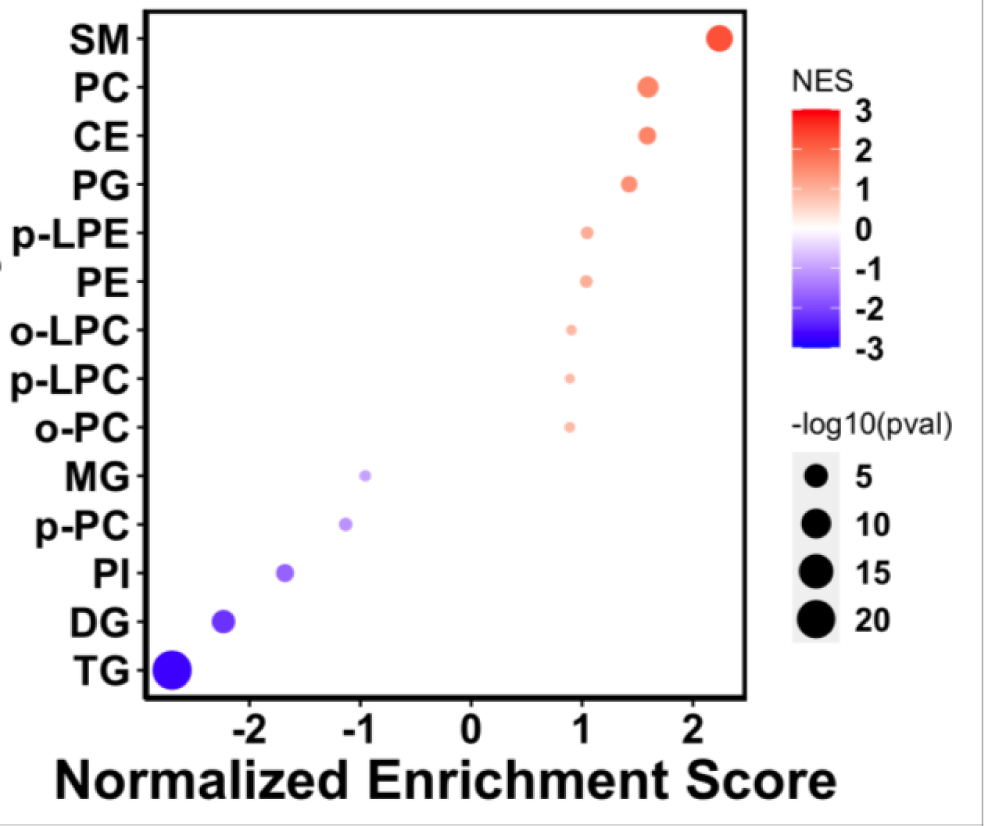
Lipid Class enrichment in adult mice under HFD. Color scale reflects the log2 transformation of the regression coefficient of the term HFD. The size of circle is -log10(pvalue).

**Figure S7B:**
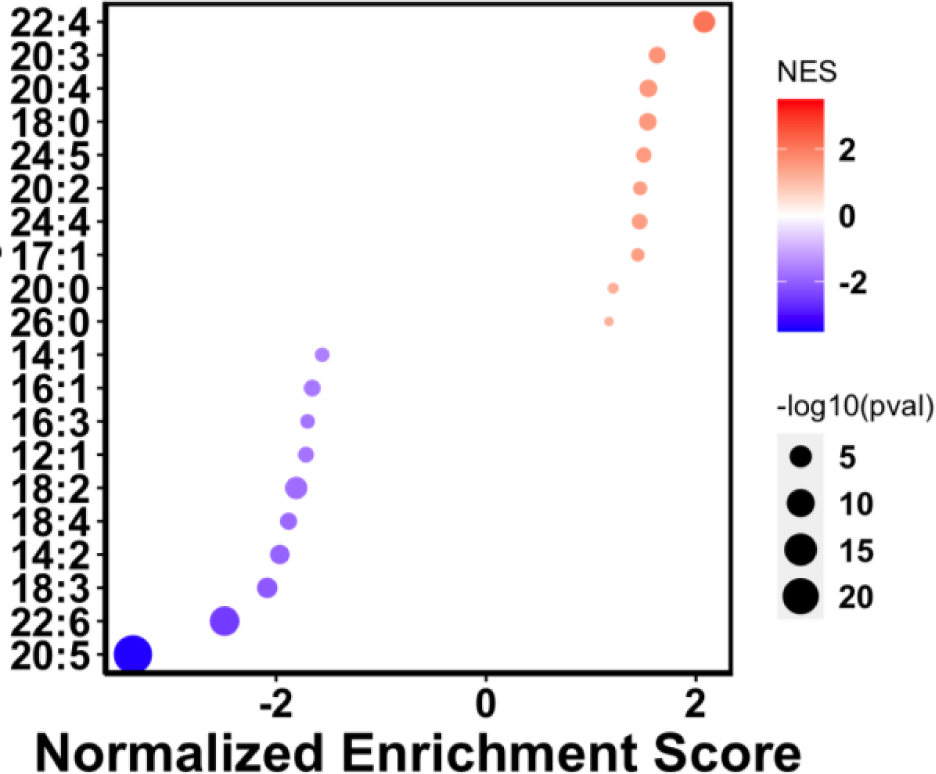
Enrichment of acyl chain composition of lipids in adult mice under HFD

